# SPARC Data Structure: Rationale and Design of a FAIR Standard for Biomedical Research Data

**DOI:** 10.1101/2021.02.10.430563

**Authors:** Anita Bandrowski, Jeffrey S. Grethe, Anna Pilko, Tom Gillespie, Gabi Pine, Bhavesh Patel, Monique Surles-Zeigler, Maryann E. Martone

## Abstract

The NIH Common Fund’s Stimulating Peripheral Activity to Relieve Conditions (SPARC) initiative is a large-scale program that seeks to accelerate the development of therapeutic devices that modulate electrical activity in nerves to improve organ function. Integral to the SPARC program are the rich anatomical and functional datasets produced by investigators across the SPARC consortium that provide key details about organ-specific circuitry, including structural and functional connectivity, mapping of cell types and molecular profiling. These datasets are provided to the research community through an open data platform, the SPARC Portal. To ensure SPARC datasets are Findable, Accessible, Interoperable and Reusable (FAIR), they are all submitted to the SPARC portal following a standard scheme established by the SPARC Curation Team, called the SPARC Data Structure (SDS). Inspired by the Brain Imaging Data Structure (BIDS), the SDS has been designed to capture the large variety of data generated by SPARC investigators who are coming from all fields of biomedical research. Here we present the rationale and design of the SDS, including a description of the SPARC curation process and the automated tools for complying with the SDS, including the SDS validator and Software to Organize Data Automatically (SODA) for SPARC. The objective is to provide detailed guidelines for anyone desiring to comply with the SDS. Since the SDS are suitable for any type of biomedical research data, it can be adopted by any group desiring to follow the FAIR data principles for managing their data, even outside of the SPARC consortium. Finally, this manuscript provides a foundational framework that can be used by any organization desiring to either adapt the SDS to suit the specific needs of their data or simply desiring to design their own FAIR data sharing scheme from scratch.

## 1. Introduction

The NIH Common Fund’s SPARC project, Stimulating Peripheral Activity to Relieve Conditions, is a large-scale program whose mission is to map the peripheral nervous system across multiple species and improve our understanding of nerve-organ interactions. SPARC achieves this aim by providing access to high-value datasets, maps, and computational studies in support of bioelectronic medicine. Bioelectronic medicine is defined as “…the convergence of molecular medicine, neuroscience, engineering and computing to develop devices to diagnose and treat diseases”^1^.

Integral to the SPARC program are the rich anatomical and functional datasets produced by investigators across the SPARC consortium that provide key details about organ-specific circuitry, including structural and functional connectivity, mapping of cell types and molecular profiling. These datasets are provided to the research community through an open data platform, the SPARC Portal available at sparc.science. SPARC is also developing new tools and technologies to support modeling and simulation of nerve-end organ interactions.

The data produced by the SPARC project is highly heterogeneous, deriving from multiple species, spatial and temporal scales, and anatomical, physiological and molecular techniques. To ensure that SPARC data adhere to the principles for making data Findable, Accessible, Interoperable and Reusable (FAIR)^2^, the SPARC curation team is charged with identifying, and implementing community standards and annotating SPARC data with rich metadata. Standards are integral to FAIR because they make it easier to combine across datasets, ensure that necessary metadata is provided, and make it possible to write automated tools to promote reuse of data. Community standards are either adopted from other domains or developed by SPARC to serve their needs.

To date, SPARC has been curating data to two primary standards developed by the SPARC consortium: 1) The Minimal Information Specification (MIS), a semantic metadata scheme capturing key experimental and dataset details; 2) The SPARC Dataset Structure (SDS), a file and metadata organizational scheme based on the Brain Imaging Data Structure (BIDS), developed by the neuroimaging community^3^. SPARC investigators are required to organize their data files and metadata according to SDS; SPARC curators then align the submitted metadata and file pointers to the MIS using automated and semi-automated workflows.

In this paper, we explain the rationale behind the design of the SDS and give a detailed description of the associated guidelines. This provides a full overview and instructions for anyone wanting to follow these FAIR data standards for any field of biomedical research. The SDS may be useful for fields where FAIR data standards are yet to be established as it is agnostic to data type. We also present automated validation and curation tools that have been developed for SPARC, which could facilitate use of the SDS beyond SPARC. The code for the SPARC curation pipelines is an open source Python project bearing the MIT license^10^. This paper also provides a foundational framework that could be used for adapting the SDS to suit the specific needs of data from a particular field of research.

## 2. Overview of SPARC Curation Process

Data and curation services and infrastructure for SPARC are provided by the SPARC Data and Resource Center. Currently, SPARC data is uploaded to the Blackfynn data platform^4^, which provides a private, password-protected space for researchers to store and organize their data. Data are uploaded from individual investigators in the SPARC consortium according to timelines and milestones negotiated with the US National Institutes of Health (NIH). Investigators are required to upload their data within 30 days of completing a particular milestone. Each batch of data uploaded to complete a milestone is considered a SPARC dataset. Investigators are given instructions and templates for organizing their data according to the SDS and are expected to upload their data in this format.

Once uploaded, data are curated by SPARC curators who will review for compliance with the SDS, completeness of data and metadata and overall quality. Certain types of data, e.g., 2D and 3D images, undergo spatial registration using the TissueMaker software developed by MBF Biosciences with organ-specific 3D scaffolds and data visualizations being created by the Auckland Bioinformatics Institute (ABI). A more detailed curation workflow is described in Section 7.

When complete, a dataset in SPARC comprises the following:

1. Data files uploaded to the Blackfynn platform organized according to the SPARC Data Structure that includes all required metadata
2. A complete detailed experimental protocol in Protocols.io describing any procedures used to obtain the data uploaded
3. If applicable, a set of fiducial mark up of 2D images for spatial registration of images to scaffolds; converting image files to required formats (performed by MBF Biosciences)
4. If applicable, data registered to 3D spatial scaffolds, which includes creating visualizations of certain types of data, e.g., RNAseq (performed by ABI)
5. A set of curator’s notes that accompanies the data file to summarize key parameters of the dataset

In this paper, we outline the rationale and structure of the SDS and some of the tooling that has been developed to support it. A separate paper will be prepared for the MIS.

## 3. Development of the SDS

To capture data across diverse types of biological data, the SPARC Consortium has adopted the Brain Imaging Data Structure (BIDS, RRID:SCR_016124) format for research objects as a foundation for the SDS (see Fig. 1). The BIDS format is a simple file folder organization and metadata scheme. At the top level, the BIDS format functions as a series of folders representing a dataset, consisting of a set of specified files and subfolders containing different types of metadata and data.

**Figure 1:**
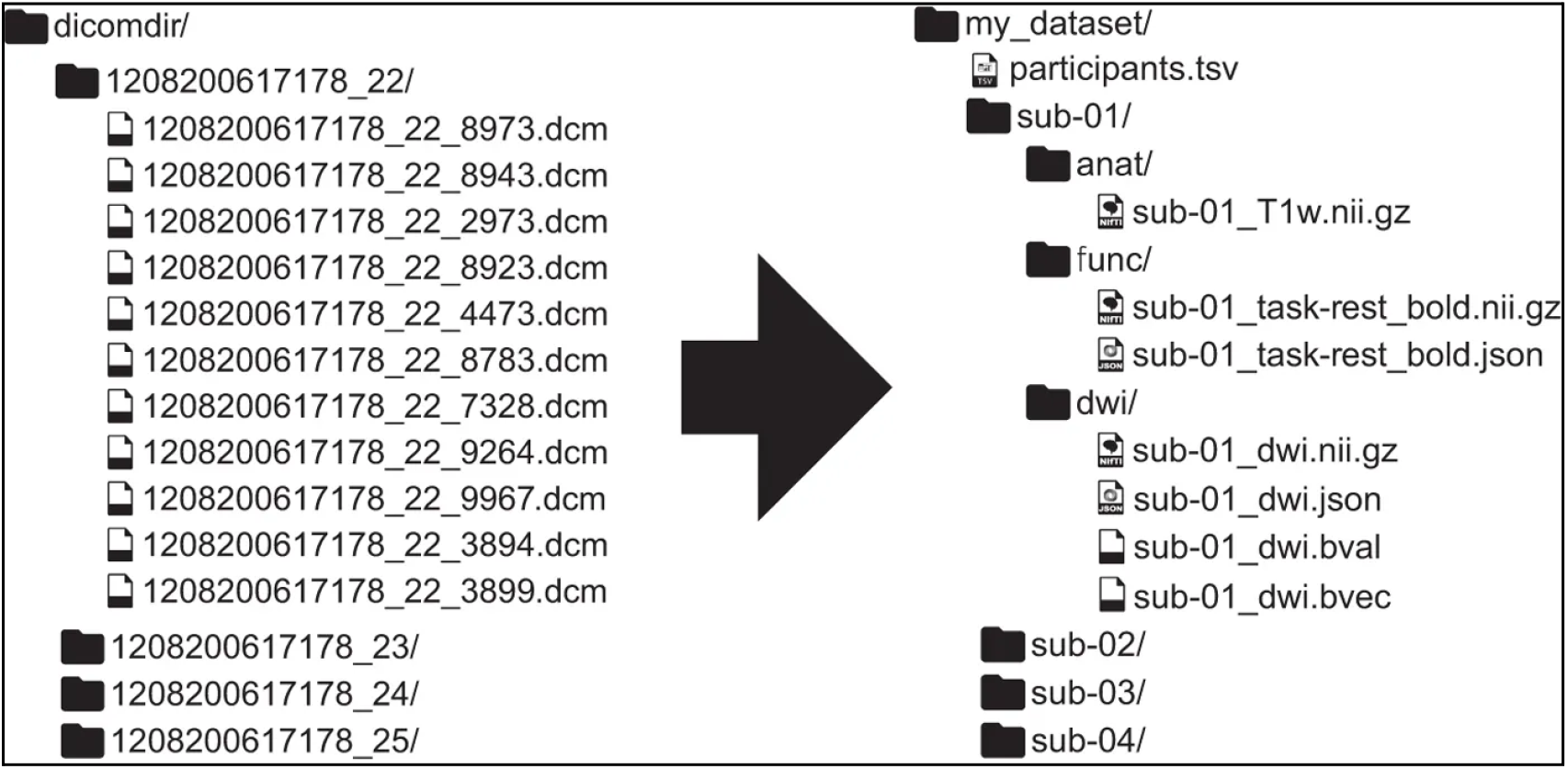
Transformation between DICOM and BIDS (Taken from Gorgolewski et al. 2016)

### 3.1. Rationale

Formal data structures, like BIDS aim to increase the integrity of scientific research through the active encouragement and facilitation of FAIR. “Findability” is improved when the names of organisms and organs are standardized to establish community ontologies. “Accessibility” and “Interoperability” are improved as files are organized in more predictable locations across different datasets and when they use common and open formats, such as csv or tiff. “Reusability” is improved by ensuring that all contributed data is well annotated and conforms to community standards, e.g., minimal information models, when such are available, and are made available under a clear license. For SPARC, all datasets are released under the CC-BY-4.0 license.

BIDS was deliberately and carefully designed to complement *likely* research practices in the laboratory to ensure accurate capture of complex imaging experiments. Towards this end, BIDS can be used by laboratories with minimal bioinformatics experience or support to manage, exchange and, submit well-annotated data in a human and machine-readable format. The BIDS format creates a resulting structure sufficiently standardized to support the creation of validation code (e.g., BIDS Validator. The BIDS validator is an application that checks for the presence of required files, and the completion of required fields within those files. The BIDS format and BIDS validation code are already used in repositories that store imaging data, including OpenNeuro.org.

BIDS was developed and refined over many years, through many meetings and by many contributors. This standard has become relatively well accepted in the neuroimaging community as a means to package and describe neuroimaging studies and has been endorsed by the International Neuroinformatics Coordinating Facility (INCF) through its standards review process^5^.

The curation team joined SPARC in 2018 just as the first deadlines for data submission by consortia members were approaching. Based on the recent INCF endorsement of BIDS, we recommended the project adopt a modification of BIDS as an initial effort to coordinate data across different laboratories. Although BIDS was developed originally for neuroimaging, its basic structure is adaptable to various experimental paradigms. Because of the diversity of data in SPARC, the large number of files and complex structure of the datasets, we felt that without a consistent structure, data in SPARC would be very difficult to work with by end users, and very difficult to curate, as each dataset would be organized and documented differently. As BIDS had already gone through multiple rounds of community review, including the independent INCF review, and is a recognized standard for the OpenNeuro Data Archive supported by the US BRAIN Initiative, we felt confident that it provided a solid foundation for SPARC in the early stages of data sharing.

### 3.2. SDS overview

The BIDS structure was modified to remove neuroimaging specific aspects, and to accommodate the fact that most data in SPARC are derived from animals and animal tissue. Thus, unlike in non-invasive neuroimaging studies, data may be acquired at the subject level, e.g., in vivo physiological recordings, or at the specimen level (from an ex-vivo tissue specimen or in vitro cell culture) (Fig. 2). The proposed modifications to BIDS were accepted by the SPARC Data Standards Committee and we moved forward with working with investigators to organize their data according to the SPARC Dataset Structure.

**Figure 2:**
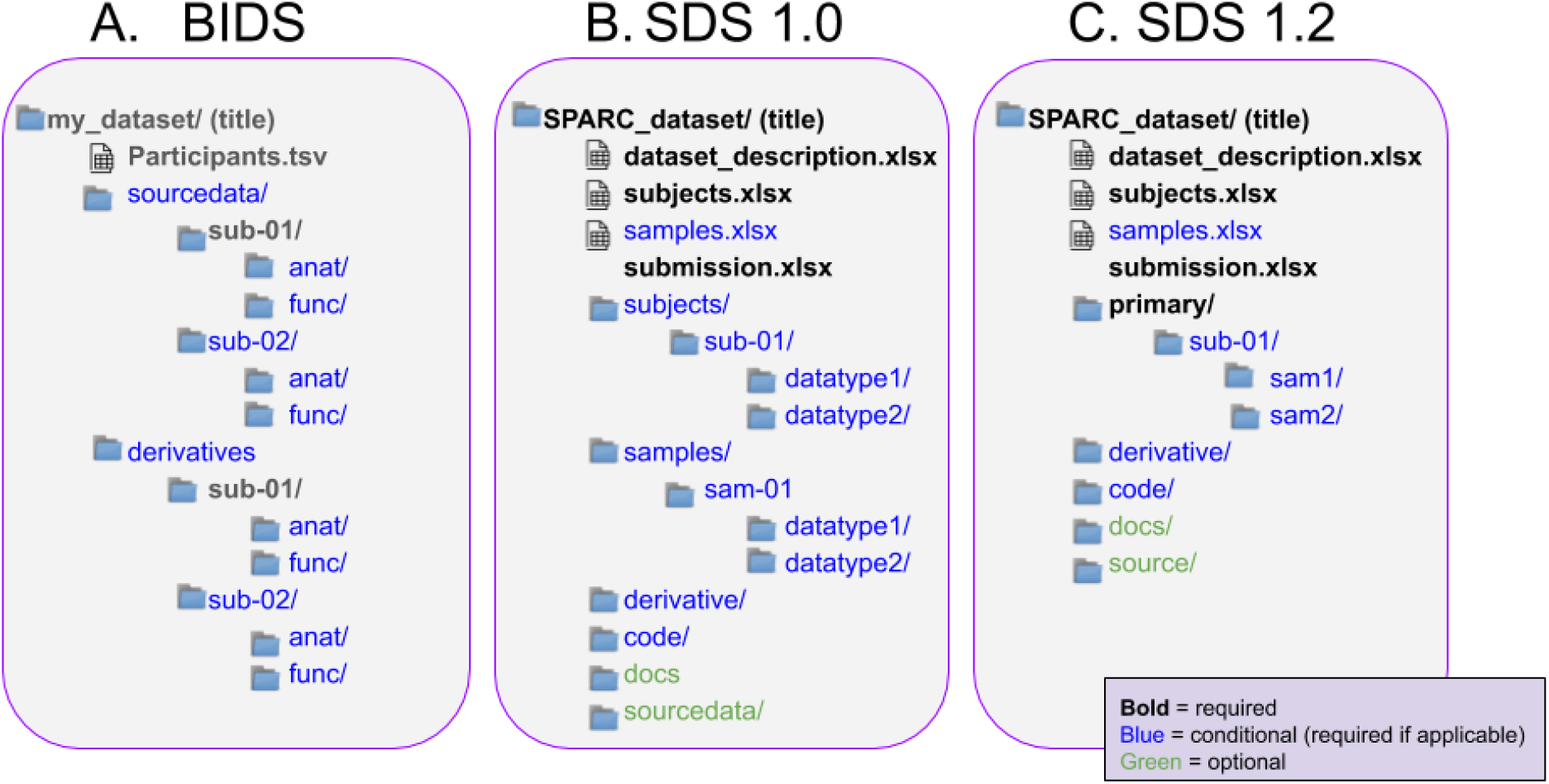
A comparison of high level details of BIDS (A), SDS 1.0 (B) and SDS 1.2 (C).

Version 1.0 was put in place to organize the first data submitted from January 2019 - July 2019 in anticipation of the debut of the SPARC data portal at the 11th Congress of the International Society for the Autonomic Neuroscience (ISAN 2019). The overall structure is shown in Fig. 2B. It defined a set of high-level folders, including one for Subjects and one for Specimens, and included various spreadsheets into which investigators could enter metadata for the dataset as a whole (dataset_description), subjects and samples. Note that the file format chosen for these spreadsheets is .xlsx, rather than an open format like .tsv or .csv. Although .csv is the preferred file format for tabular data in SPARC, the curation team wanted to make it easier for both investigators and curators by including features such as drop down value sets for certain metadata fields, features which are not supported by these basic formats. In addition, the Blackfynn data platform did not have a viewer available for .csv files, but did support on-line viewing of .xlsx through the Microsoft Open Office suite. As with BIDS, the SDS follows the inheritance principle that requires any metadata files in the root directory to apply to all folders and files below it, except when explicitly overridden by a metadata file contained in a lower order folder.

After a review of datasets submitted for ISAN and interviews with investigators, the curation team modified the basic structure to simplify the folder structure (Fig. 2C), collapsing the subject and samples folders into a single folder named primary. Samples may now be nested under their respective subjects. The current release is version 1.2, (Fig. 2C). The required folders and files are provided to investigators as a downloadable versioned template via GitHub (https://github.com/SciCrunch/sparc-curation/releases/tag/dataset-template-1.2.3). All datasets are now curated according to version 1.2, including those that were released for the ISAN 2019 meeting.

Although the SDS is modeled on the approach by BIDS, i.e., file folder organization, naming scheme and provision of critical metadata, it is sufficiently distinct from BIDS that we do not consider it an extension, but rather a derivative (see Fig. 2 for a comparison between BIDS and SDS). We now describe SDS V1.2 in more detail.

## 4. SPARC Data Structure V1.2

The SPARC Dataset Structure includes the following components (Fig. 3):

- A set of organized data files in a hierarchical set of predictably named folders and subfolders. Folders/subfolders may contain supplementary and additional documentation, i.e. manifest files that describe the files and/or folders contained therein.
- A set of descriptive top-level files that contain information on subjects, experimental information, and dataset descriptions. These descriptive files include both spreadsheets containing structured metadata and text files with additional information.
- A set of file manifests associated with each folder that provides descriptions of the contents.

**Figure 3:**
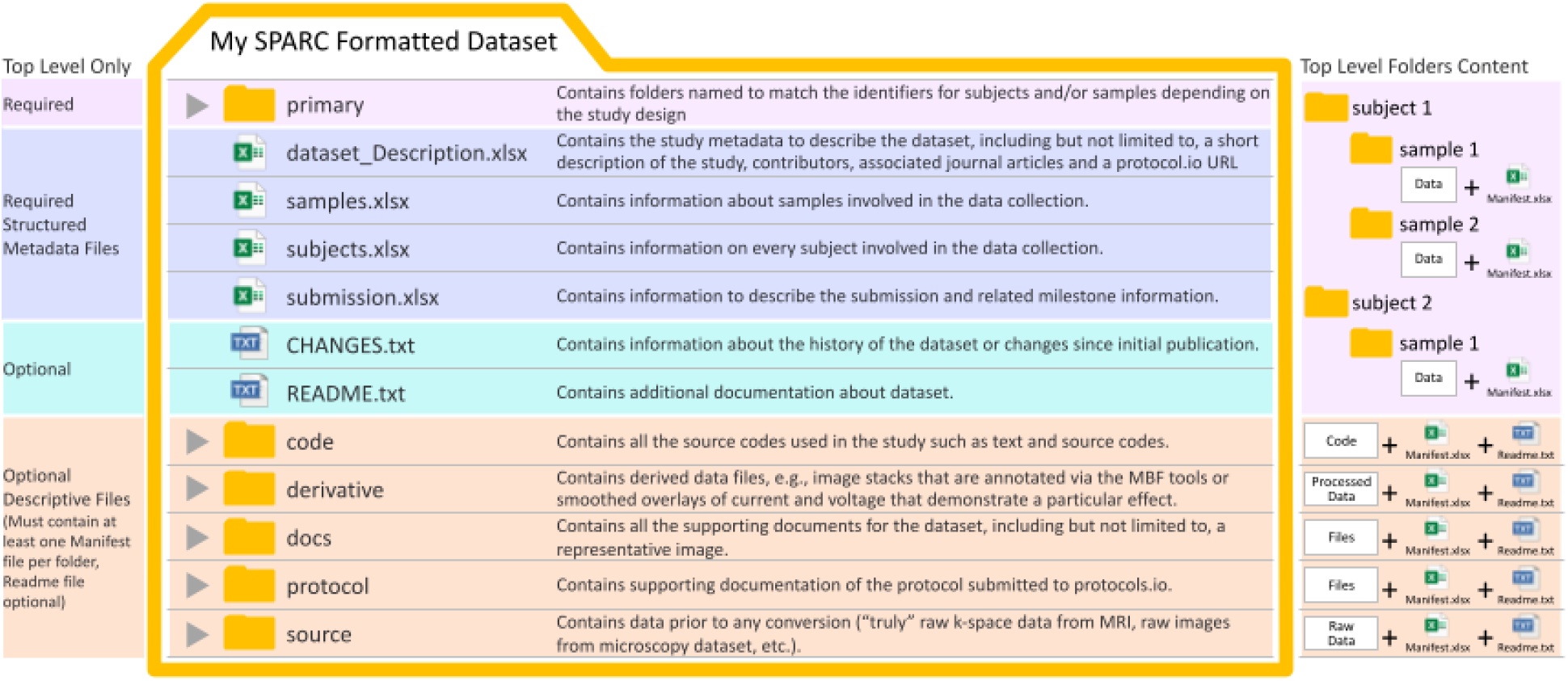
The organization structure of the files and folders for a SPARC dataset.

### 4.1. Top-level structure

Data files are organized into 3 different top-level folders, depending on the type of data:

- **primary**: a required dataset dependent folder that contains all folders and files for experimental subjects and/or samples, e.g., time-series data, tabular data, clinical imaging data, genomic, metabolomic, microscopy data. The data generally have been minimally processed so they are in a form ready for analysis. Within the primary folder, data is organized by subjects or samples (see Section 5). All subjects and samples will have a unique folder with a standardized name corresponding to the exact names or IDs as referenced in the subjects and samples metadata file (see Fig. 3).
- **source**: an optional folder containing unaltered, raw files from an experiment, if they are included in the data. For example, this folder may include the “truly” raw k-space data for a Magnetic Resonance (MR) image that has not yet been reconstructed, or a set of microscopic images that had not yet been assembled into a mosaic. The reconstructed DICOM or NIFTI files and the image mosaic, for example, would be found within the primary folder.
- **derivative**: a required folder if derivative data exists. This folder contains derived data files. For example, processed image stacks that are annotated via the MicroBrightField (MBF Biosciences) tools, segmentation files, or smoothed overlays of current and voltage that demonstrate a particular effect. If files are converted into a format other than what was submitted, these files are included in the derivative folder. Derived data should be organized into subject and sample folders, using the subject and sample IDs as the folder names, as with the primary data. Other files are organized in three different (optional) folders:
- **code**: a required folder only if code is used in generation of the data; the folder contains all the source code used in the study, e.g, MATLAB.
- **protocol**: an optional folder that contains supplementary files to accompany the experimental protocols submitted to Protocols.io. The additional files in this folder are not a substitution for the experimental protocol which should have been submitted to Protocols.io/sparc.
- **docs**: an optional folder that contains all the supporting documents for the dataset, including but not limited to, a representative image for the dataset. Unlike the readme file, which is necessarily a text document, docs can contain documents in multiple formats, including images.

### 4.2. Descriptive top-level files

A set of descriptive, top-level files contain information on subjects, samples, dataset descriptions and administrative data. These files contain required metadata fields that are aligned to the DataCite schema (dataset description), and the HBP’s (Minimal Information about a Neuroscience Data Set)^6^ for subjects and samples. Additional recommended fields are included for each (see Supplementary Material A). Investigators are encouraged to add additional columns beyond this core set to thoroughly describe the dataset.

While there is a great deal of flexibility built into the metadata templates in order to accommodate the diversity of experimental paradigms and data, for the effective functioning of the validator (described in Section 6), it is important for data wranglers not add, edit or delete required columns in the mandatory descriptive files (these are color-coded (see green and blue in Appendix A). If there is information that doesn’t correspond with available columns, the information should be added to a new column on the right-hand side (subjects and samples) or a new row on the bottom of the sheet (dataset description). If there is information not available to the researcher at the time of submission, fields should be left empty or marked “unknown”.

An overview of the spreadsheet metadata templates is provided below:

- **dataset_description** (xlsx, csv or json): Required file containing basic metadata about a dataset, derived largely from the DataCite Schema^7^. A full list of metadata and definitions is provided in Supplementary Table A1. Investigators provide basic metadata such as title, description, contributors, funding and contact person, that provide provenance for the dataset and also support formal data citation. The version 1.2 release includes an additional field specifying the metadata version. This field is <u>not to be changed</u> by data submitters. It allows proper alignment between different metadata releases, securing the data integrity for multiple batches of submissions. We also encourage researchers to describe if they plan to submit more data for new or for the same subjects, i.e., this dataset is part of a larger study. This will help determine when all the primary data has been deposited and help with mapping across the different parts of the dataset.
- **submission** (xlsx, csv or json): Required file containing information relevant to internal SPARC bookkeeping, relating milestones negotiated with NIH to datasets submitted. According to the SPARC Material Sharing Policy^8^, data is to be deposited within 30 days of milestone completion and will become public no later than 1 year after milestone completion. This file is for internal use only; it must not be released when the data are published.
- **subjects** (xlsx, csv or json): Required file if subjects are used in the experiment producing the dataset. Contains updated fields with required and optional metadata fields providing information about subjects (model organism or animals) involved in data collection. The file contains fields specifying provenance for the subject, e.g., subject_id, pool_id and experimental group (blue fields in Appendix A2). Each subject and pooled subjects must be assigned a unique ID, as this ID is used to name the data folders for individual subjects. For proper mapping of the data, folders containing experimental data need to <u>exactly</u> match the subject ID. All subject identifiers must be unique within a dataset and not contain any sensitive, identifiable information (for human subjects). Having each lab use unique subject identifiers across datasets is highly desirable to aid in connecting multiple experiments using the same subjects. In the future, we plan to connect subjects across datasets and projects; however, we currently do not map subjects across multiple data submissions. The subjects.xlsx file contains several mandatory fields (green in Appendix A1) including species, age, strain and Research Resource Identifier (RRID). Additional columns containing additional descriptive metadata, demographic assessment data, etc, largely derived from OpenMINDS, are provided for investigators in the template. In the download template, these are highlighted in yellow (See Appendix A1) and serve as exemplars of the types of metadata that are important for providing scientific context. According to the FAIR principles, data should be described by a “plurality of relevant attributes”, but we are leaving it up to the investigators’ discretion to decide what is sufficient for others to understand and reuse the data. Investigators have the liberty to add as many fields as needed that they deem necessary. Currently, all metadata provided for subjects and samples is provided in free text, which is then mapped to the SPARC vocabularies by the curation team (see Section 6.1). However, we are actively working with investigators on lists of controlled vocabularies for certain fields.
- **samples** (xlsx, csv or json): Conditional file required if measurements are obtained from samples, e.g., tissue slices, derived from individual or pooled subjects. This file contains information about samples used to generate the data. Investigators must provide a unique ID for each sample that will be used to name the data folders. The sample ID must match the folder ID exactly. Each sample should also reference a subject from the subject file; a single subject (a research animal/donor) may be linked to multiple biological samples derived from that subject. If the samples are pooled from multiple subjects, the complete provenance must be specified in the subject file. The metadata present in the samples file should also explicitly note whether a sample was collected directly or was derived from another sample. Required metadata includes the subject or tissue from which the sample was derived and the anatomical location (green in Appendix A3) Additional Fields may be added by the investigator. The template provides some suggested fields derived from the Minimal Information about a Neuroscience Dataset (OpenMINDS). *Investigators should only use columns that are relevant to their type of study*.

An overview of the descriptive text files is provided below:

- **README** (txt): Required file provided by Investigators that contains necessary details for reuse of the data, beyond that which is captured in structured metadata. Some information that should be included are:

- How would a user use the files that are provided? e.g., first open file X and then look at file Y.
- What additional details do they need to know? Are some subjects missing data?
- Are there warnings about how to use the data or code?
- Are there appropriate/inappropriate uses for this data?
- Are there other places that users can go for more information? e.g., did you provide a GitHub repository or are there additional papers beyond what was provided in the metadata form?
- **Plog** (xlsx or txt): Optional performance log file, which can be used to attach information about individual performances of the experiment, e.g. how long they took, what the average room temperature was, or who performed them. There is currently no other place in the data model to attach that kind of information.
- **CHANGES** (txt): Conditional file required if a new version of the dataset is uploaded to document any changes from the previous **version.**

### 4.3. Manifests

Manifest (xlsx, csv or json): Required file that must be in all folders containing data files (Fig. 3, Fig. 4)^9^ and in folders with subfolders whose meaning is not clear. This file contains information and metadata about the files and folders that are expected in the folder where they sit. Required fields include file name (or file name pattern for folders with many related files), description and file type, although investigators have a lot of flexibility by adding additional columns, including notes about pertinent aspects of each file that differentiate the files (e.g., data collection specific protocol, stimulation condition, microscope filter applied, drug applied, etc). The manifest file can apply to collections of files (through the use of a file pattern) or list specific files (e.g. sub??-task1-run?? can specify all the files related to task1 in the protocol). If investigators include folders that organize data along a particular dimension, e.g., datatype or time point, a manifest file should be generated that describes the content of the folders.

**Figure 4.**
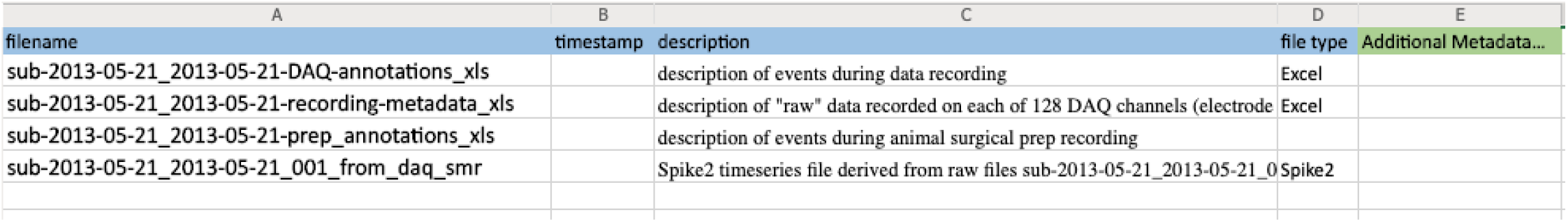
Example of a complete manifest. From Morris et al (2020).

### 5. Folder hierarchy principles

The folders and files pictured in Fig. 3 are required and invariant for each SPARC dataset. This invariance imposes a standard structure for SPARC datasets that allows a user to reliably navigate the often complex experimental data (Fig. 5).

**Figure 5.**
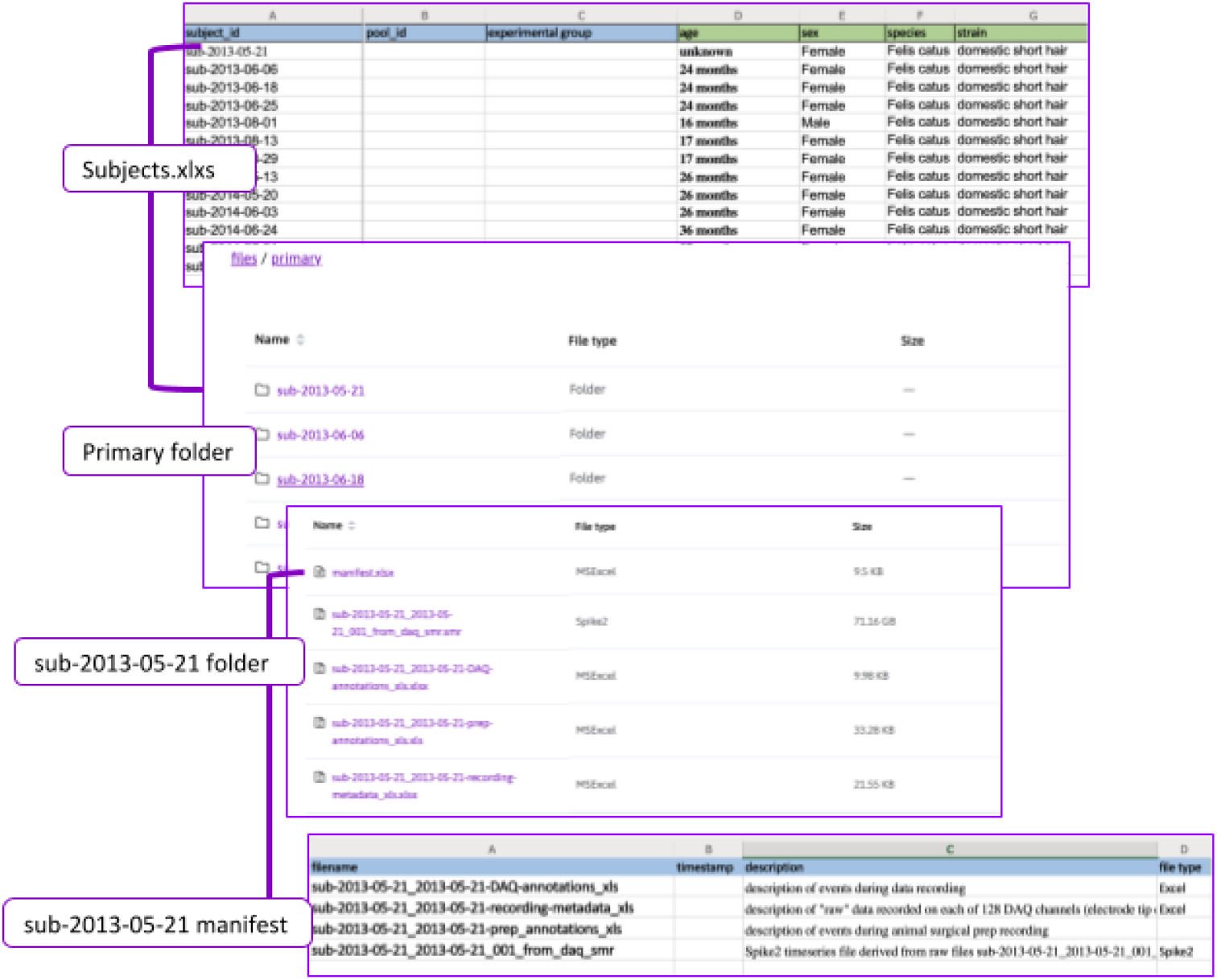
Relationships between metadata files and folder structure. Example taken from (Morris et al. 2020).

However, given the variety of different experimental protocols and the way in which subjects and samples are treated across different types of experiments, the folder and file structure can vary among different datasets within the primary data folder. Some examples are provided with the template download (Fig. 6).

**Figure 6:**
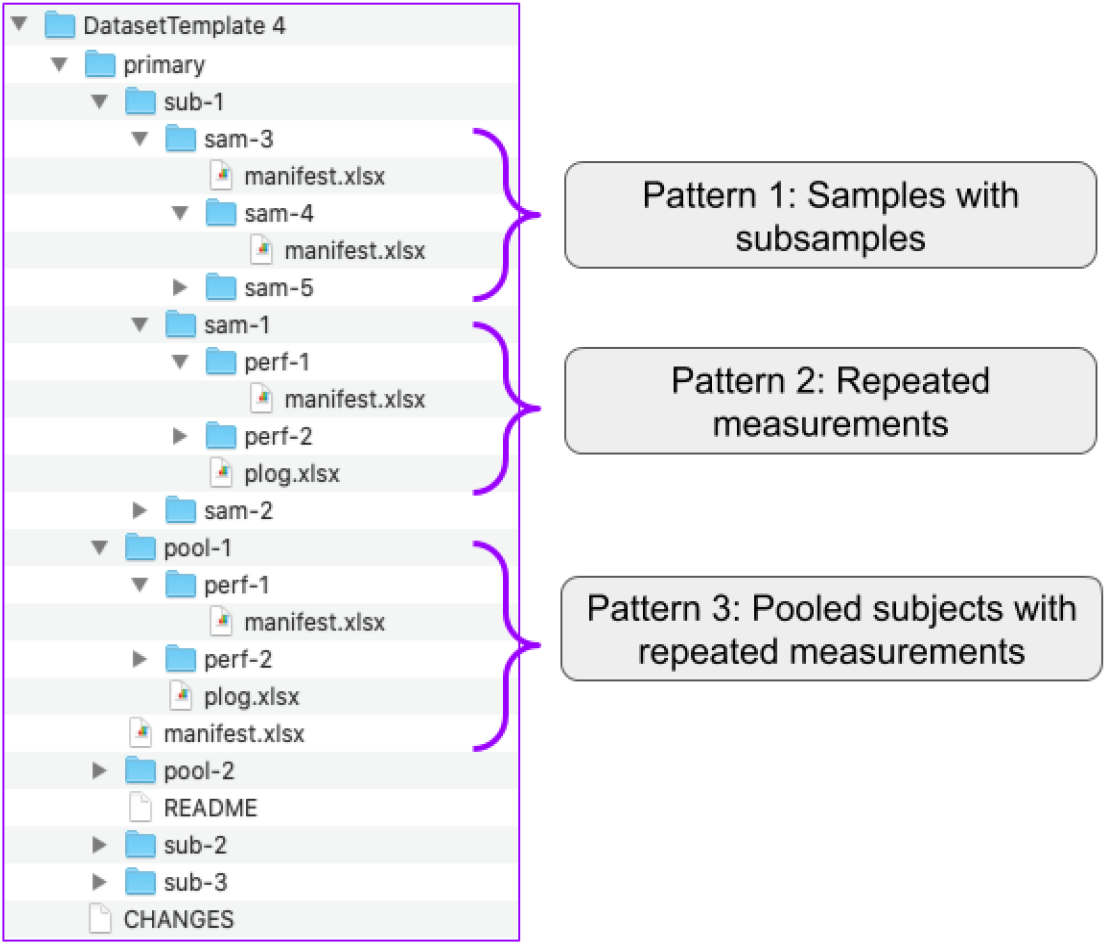
Dataset-template 1.2.3 folder hierarchy.

For the majority of SPARC datasets, data in the primary data folder are organized into subject folders, with the folder names corresponding to the subject IDs provided in the subjects.xlsx file. If samples are derived from these subjects, data files are organized within sample subfolders under the appropriate subject, according to this pattern (Pattern 1)

The inheritance principle applies, so that if sample 3 (sam-3) appears as a subfolder of subject 1 (sub-1), then it is assumed that sample 3 was obtained from subject 1 (Fig. 5). In some cases, no data may be derived from the subjects directly, i.e., no data files are generated at the subject level. In this case, investigators could omit the subject folder (although the subject.xlsx file must be included to provide the appropriate metadata).

### Time series data

For functional studies where measurements are obtained at different time points, different time points should be organized into folders labeled perf-1, perf-2 etc, where the numbering indicates the temporal ordering, under either the sample or subject folder. An example is shown in Fig. 6 (Pattern 2). Note that in this case the manifest file would specify information about the data that will be found in the perf-1 and perf-2 folders.

### Pooled samples or subjects

Although the majority of data are organized with subjects nested under the primary data folder and samples nested under subjects, this simple hierarchical arrangement does not apply to all datasets. In some cases, samples may be pooled from multiple subjects, in which case the sample folder lives alongside the subject folder and not nested within it, according to Pattern 3. The SDS also accommodates subject pools where the samples folder is replaced with the **pooled** folder (Pattern 3; Fig. 6).

Note that pool_ids and characteristics must be provided in the subjects.xlsx file.

## 6. Tooling to support SPARC Dataset Structure

### 6.1. SDS Validator

To enforce the SDS structure and required metadata fields, the curation team developed a SPARC Dataset Structure validator^10^ that is used for frequent checks to ensure the integrity of the data across the platform and provide valuable feedback to the curation team. The validator is written in python and uses JSON schema^11^ to specify the expected structure of the dataset files and folders, as well as the structure and contents of the 4 types of metadata files (dataset_description, subjects/samples, submission, and manifest). Tabular metadata files are transformed into JSON, and validated against the schemas.

The validator first checks that all the required metadata files are present after which the content of the individual metadata files is validated. For example, in the subjects.xlsx file, checks are performed to ensure that all subject ids are unique, that there are not names in columns that expect numbers (e.g. ‘adult’ in the ‘age’ column is an error) and that the files in the primary data folder match the names and number of subjects and samples provided in the metadata files. The validator also checks that organism and anatomical entities are present in the appropriate columns, by matching the content of these columns against the SPARC vocabularies (see section 6.1). This is not an exhaustive list of the checks that are performed, but it gives a flavor for the types of checks that are done (Fig. 7).

**Figure 7:**
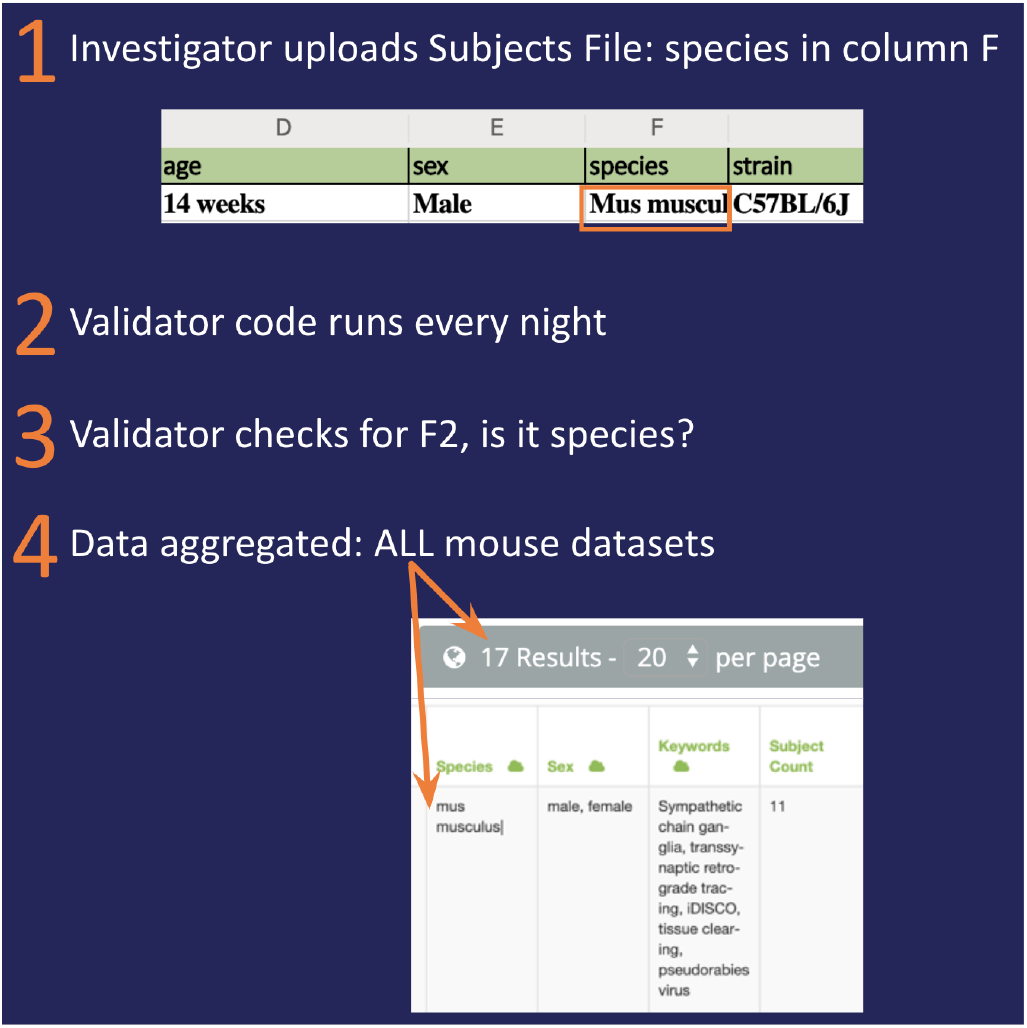
Workflow for the SDS Data validator

Some of the most common mistakes detected by the validator arise when investigators remove headers or cells for which they do not have information. This means that the validator looks for information in the wrong place, e.g. if species information is expected in cell F2 (Fig. 5), but the investigator deleted column E, then the strain information will be noted in the species field producing an error because c57BL/6J is not in the ontology as a valid species name, and all other information to the right of the deleted column or below the deleted row will also be incorrect.

Errors are noted per dataset and categorized by type for curation so that curators can act on the error. For simple alignment errors, the curators usually replace the affected files by pasting misaligned data into a fresh template. With the newly released data organization tool, SODA, (see Section 6.2) these sorts of errors will be less of a problem because at least some of the metadata files will be replaced by a form that asks investigators questions and produces a properly formatted file.

The process of validation is done automatically on each dataset, but is only meaningful for datasets that are undergoing curation, where these errors are read and acted upon. While the data are being prepared by the investigator for submission, it is not uncommon for datasets to have very large error numbers as none of the files may be in the right location and metadata fields may be incomplete. The complete curation workflow is described in Section 7.

In addition to running the validation of the required metadata, the validation code also extracts metadata from the SDS, and maps them to the MIS. During this process, certain metadata fields, e.g., anatomical structure, are mapped to the NIF Standard Ontology^12^ (RRID:SCR_005414), which in turn imports multiple community ontologies such as NCBI Taxonomy, UBERON, ChEBI. Additional ontologies, e.g., FMA are used as necessary. A list of identifiers used to map SPARC data is provided in Table 1.

**Table 1:**
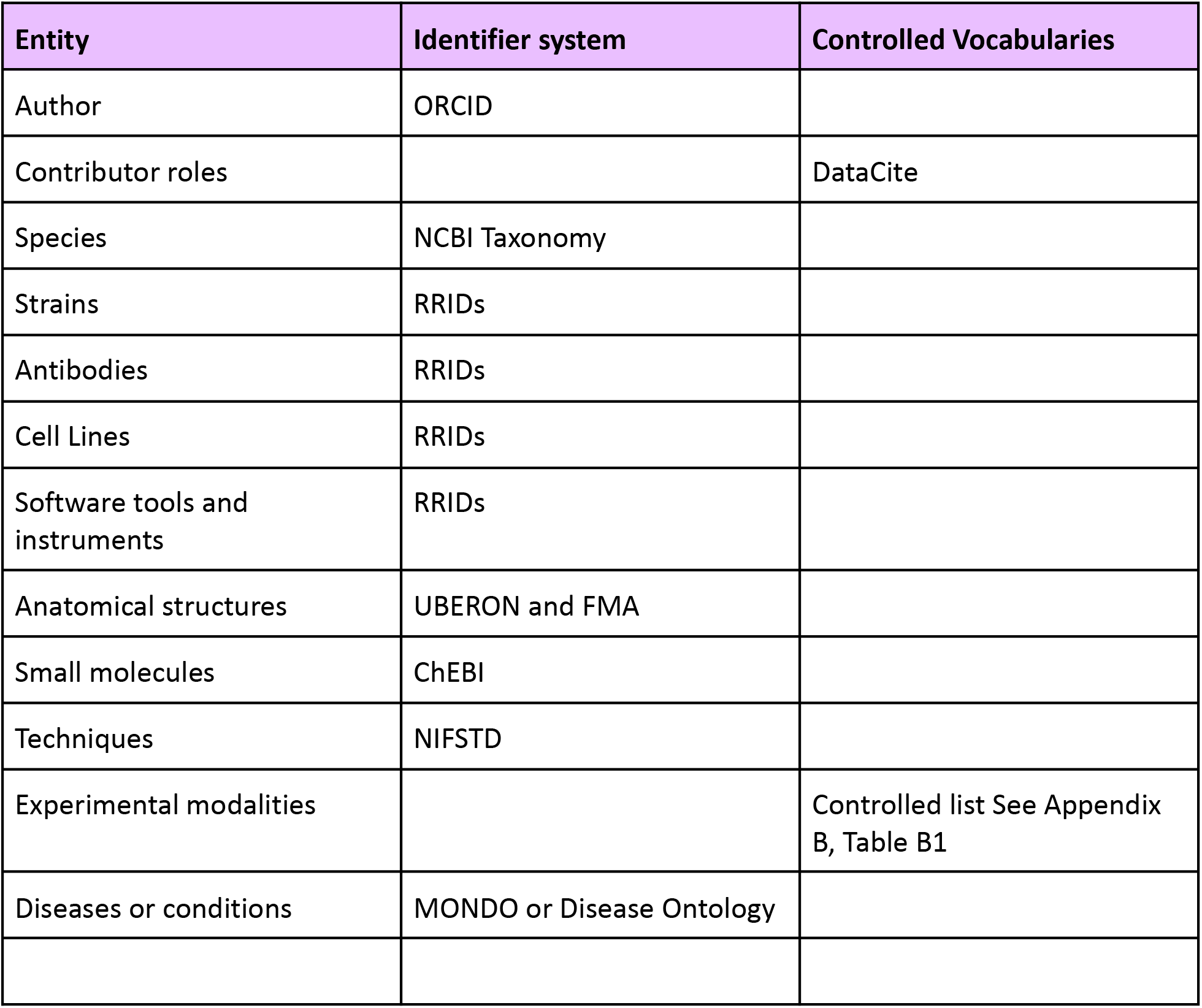
List of ontologies and controlled vocabularies used to map SPARC metadata.

| Entity | Identifier system | Controlled Vocabularies |
| --- | --- | --- |
| Author | ORCID |  |
| Contributor roles |  | DataCite |
| Species | NCBI Taxonomy |  |
| Strains | RRIDs |  |
| Antibodies | RRIDs |  |
| Cell Lines | RRIDs |  |
| Software tools and instruments | RRIDs |  |
| Anatomical structures | UBERON and FMA |  |
| Small molecules | ChEBI |  |
| Techniques | NIFSTD |  |
| Experimental modalities |  | Controlled list See Appendix B, Table B1 |
| Diseases or conditions | MONDO or Disease Ontology |  |

The validator produces a set of files from the contents of the required metadata files, the ontologies and other data sources such as protocols.io. These are made available in several formats including the MIS “ttl” file (also JSON and CSV), to Blackfynn, curation systems, and the DRC staff. With each run of the validator code the metadata in the ttl file will therefore change to reflect the current state of the dataset. The curation team has created a private searchable and sortable table using UCSD’s SciCrunch.org infrastructure, https://scicrunch.org/sparc, which allows curators to quickly see which elements are missing in each dataset and determine if the error can be fixed by curators or whether the investigator needs to help resolve issues.

### 6.2. Software to Organize Data Automatically (SODA) for SPARC

Complying with the above-described guidelines requires additional time investment from the researchers (as with any data curation and submission standards), and the data curation process can become progressively overwhelming as additional data is submitted. If researchers are not currently using any standard way of organizing their data within the laboratory, in the long run, this work will benefit the laboratory. However, if researchers already use a formal method for organizing their data, complying with SPARC requirements could prove even more burdensome as they must organize their data according to additional rules. To remediate this issue, a software named Software to Organize Data Automatically (SODA) for SPARC has been developed to assist SPARC investigators in easily curating and annotating their datasets.

Distributed as an open-source (MIT license) and cross-platform (Windows, macOS, Linux) desktop application, the goal of SODA for SPARC is to bridge a long-standing, overlooked gap between comprehensive data standards and their convenient application by researchers. SODA for SPARC provides an interactive interface that, without requiring any coding knowledge, walks SPARC investigators step-by-step through the SPARC data curation process, all the while automating repetitive, complex, and time-consuming tasks. Besides being time-efficient, SODA for SPARC also provides the convenience to SPARC investigators of organizing their datasets following a custom workflow (e.g., based on personal preferences or to comply with internal guidelines applicable in their labs) and rapidly organize their data according to the SDS only when they are ready to submit the dataset for review by the SPARC Curation Team.

The SODA for SPARC installers as well as the source code are accessible via the dedicated GitHub repository^13^. During the first phase of development (May 2019-August 2020), the following features were integrated into SODA for SPARC (Fig. 8):

1. Prepare submission and dataset_description metadata files through an intuitive interface and with assistance from the program that provides access to standard values/terminologies and makes automated suggestions based on previously saved information.
2. Prepare datasets step-by-step via a convenient interface:

- Specify desired local data files to be included in each of the SPARC folders.
- Specify metadata files to be attached.
- Request manifest files to be generated automatically.
- Check that information provided during the previous steps will generate a SPARC-approved dataset using an automated validator (before a thorough validation by the SPARC Curation Team).
- Generate a dataset based on information specified during the previous steps either locally or directly on the Blackfynn platform (to avoid duplicating files on the user’s computer).
3. Manage datasets by easily connecting to Blackfynn with SODA for SPARC then conveniently create datasets, add metadata to Blackfynn datasets, manage dataset permissions, upload local files/folders, and share datasets with the SPARC Curation Team for review.

**Figure 8.**
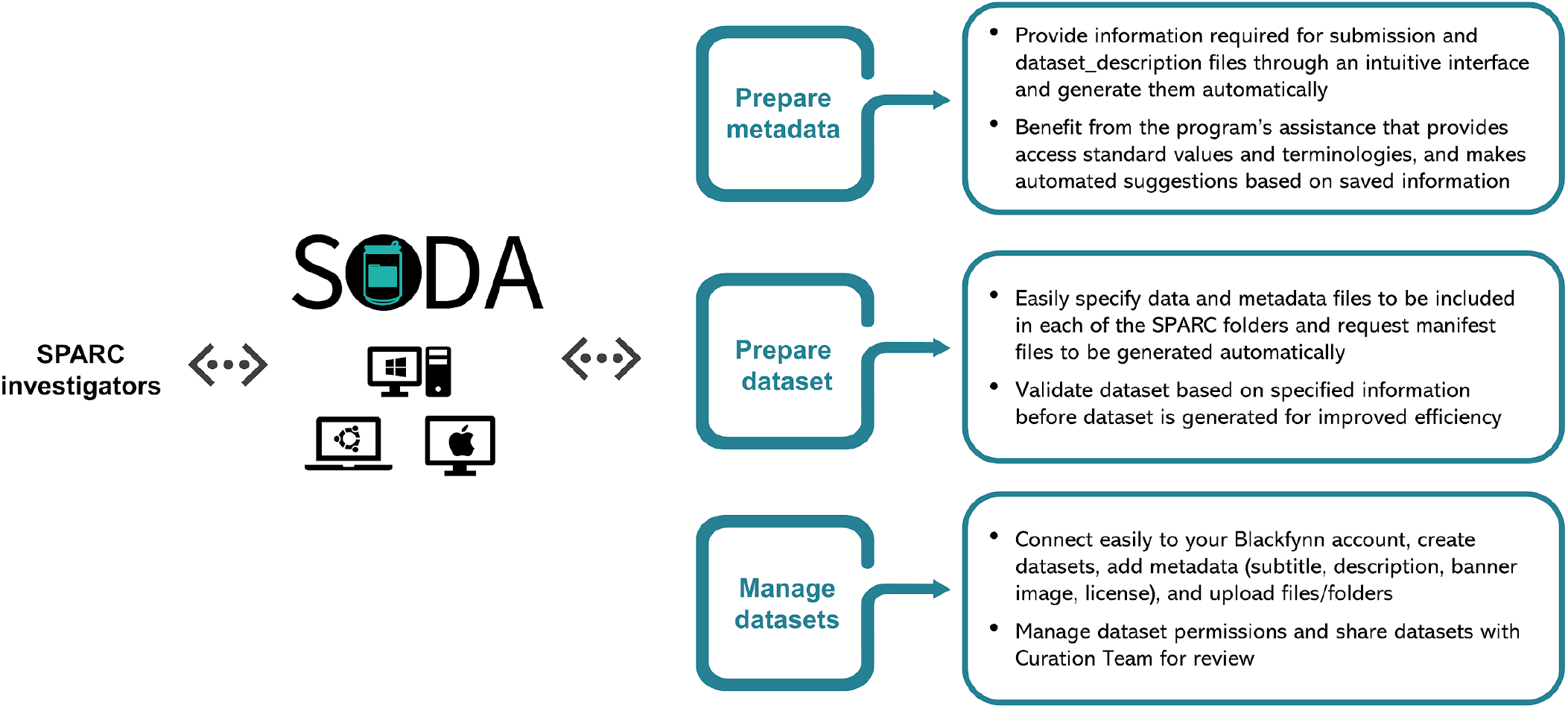
Overview of the major features included in SODA during the first development phase.

During the second phase of development (starting September 2020), more features are being added to the software including a virtual interface for organizing data, support for collaborative data curation, assistance for preparing samples and subjects metadata files, and file-level curation support. The user interface is also being upgraded to make use of the software more intuitive. A screenshot of the user interface from the current version (3.0.1) is provided in Fig. 9. A team of 10 beta testers, all of whom receive funding from the SPARC program, is reviewing and providing feedback frequently to ensure that SODA for SPARC meets the needs of the SPARC investigators. Preliminary testing by the beta testers has shown that computer-assisted curation with SODA not only reduces the time required by investigators to organize and submit their data, but also minimizes human errors^14^.

**Figure 9.**
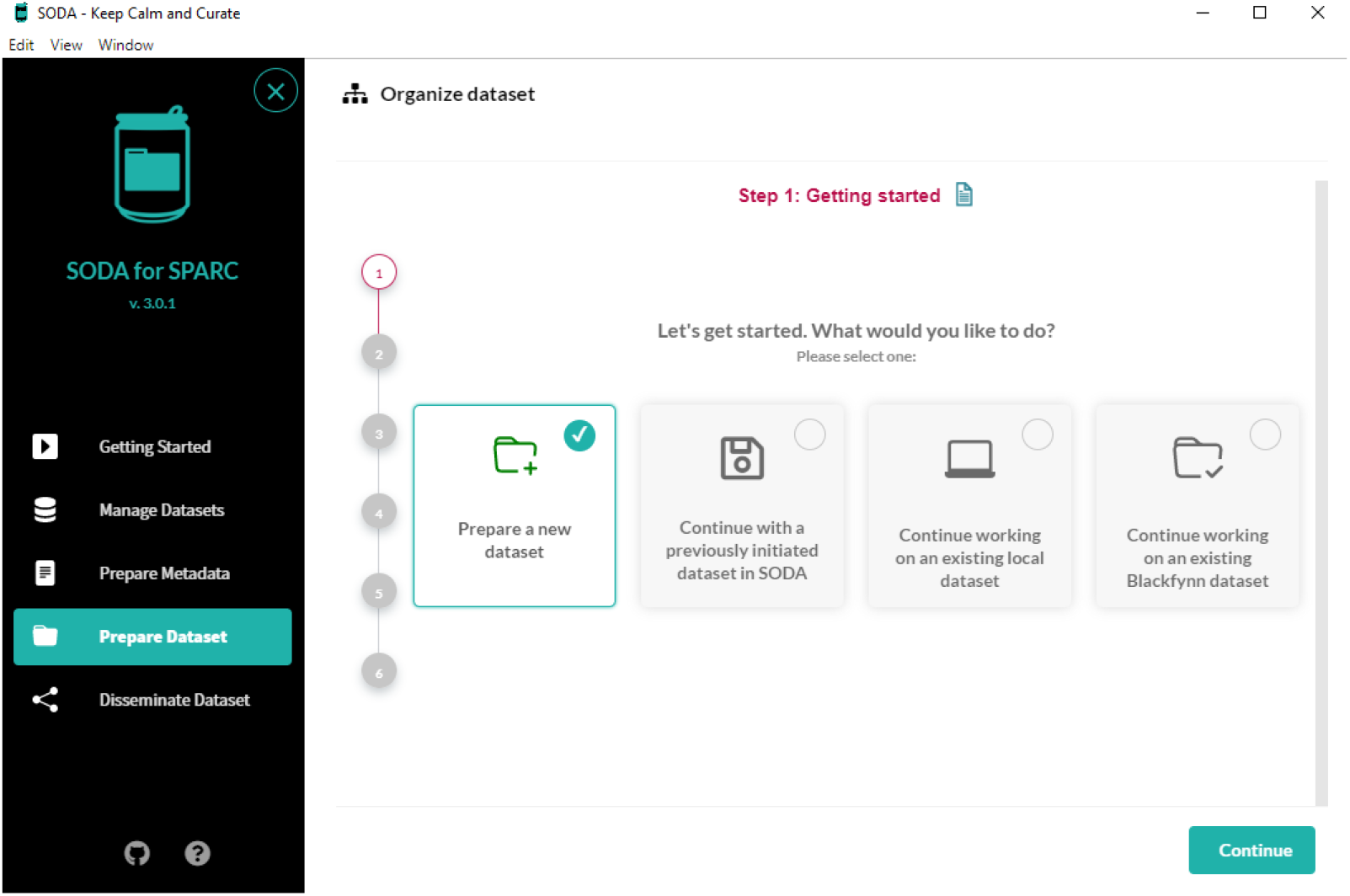
User interface (on a Windows computer) from version 3.0.1 of SODA for SPARC.

More features will be included in the future to enhance further the curation workflow and ensure that SPARC datasets are disseminated efficiently. Even beyond the SPARC consortium, quality data curation is a critical concern. SODA for SPARC could impact the broader research community by providing an exemplar, foundational tool for convenient and time-efficient data curation, which could then be adopted by other projects. In the future, we expect to modify the BIDS-inspired SPARC SDS for computational studies (the changes as they currently stand are in a draft version and will need to be approved by the data sharing committee before being acted on) that are undertaken as part of the SPARC project, it is likely that this will involve changes to the SODA for SPARC tool in compliance.

## 7. SPARC data submission workflow

All Investigators in SPARC have 1 year from the time a milestone is completed (Fig. 10), and a draft dataset is submitted (step 1) to publish the resulting dataset (step 4). A dataset is published when it has been assigned a digital object identifier (DOI) and is available for viewing and download by the public. During that year, the dataset will move through several curatorial stages and possibly an embargo period. Investigators will have 30 days from the completion of a milestone to formally submit their data to the SPARC Data Repository. Data is considered completely submitted only when the data are shared with the Data Curation Team. Once curation is complete, the dataset moves into an embargo phase or is published. During the embargo phase, the data set is visible only to members of the SPARC consortium who have signed a data use agreement. The submission + curation + embargo period add up to 1 year, that is, the length of the embargo period depends on how long it takes to curate the data to the above standards. Curation is a collaborative process that involves a back and forth between the investigator and the curation team and so the time to completion is difficult to predict. However, if investigators wish to publish before the end of the embargo period, they are encouraged to do so.

**Figure 10:**
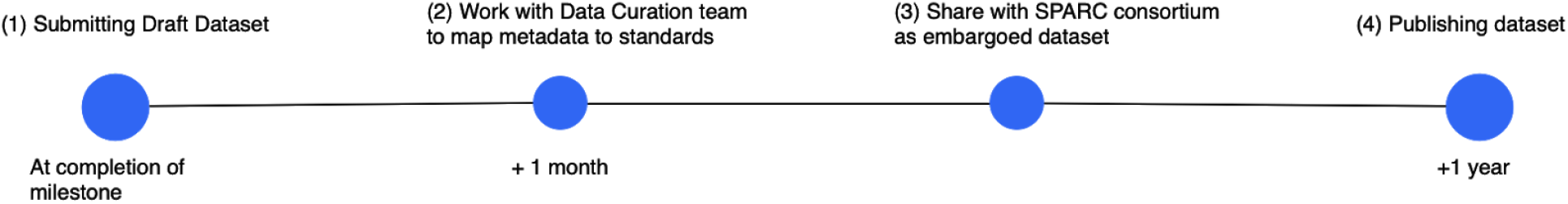
Data submission milestones

Creating a SPARC dataset in the SDS structure involves multiple steps. Instructions for creating a dataset with detailed steps can be found at https://sparc.science/help/7k8nEPuw3FjOq2HuS8OVsd):

- INVESTIGATOR: Create and name a draft dataset in their private space on the SPARC Data Repository hosted by Blackfynn (within the “SPARC Consortium” organization on Blackfynn).
- INVESTIGATOR: Organize and upload files to the dataset within this space according to the requirements of the SPARC Data Structure, using the template provided by SPARC.
- INVESTIGATOR: Request a publication review. This step initiates the curation process and locks the dataset so that changes can only be made by the curation team
- INVESTIGATOR: Upload the experimental protocol to Protocols.io and share this protocol with the SPARC group.
- SCRIPT: Downloads all SPARC data from Blackfynn.
- CURATOR: Logs all SPARC datasets into the master spreadsheet, their status and any communication tickets with the investigator.
- SCRIPT: Run weekly to find new datasets by matching the dataset IDs in the data dump with those on the master spreadsheet.
- CURATOR: Send an email acknowledgment when new dataset is detected within 5 working days.
- SCRIPT: Run all datasets through the validator
- CURATOR and INVESTIGATOR: Curators will work with the automated validator report and investigators to ensure that required fields are complete and the folder structure is appropriate.
- CURATOR: Find MIS data elements in the protocol using semi-automated tools, adding these to the structured metadata package that will be sent back to Blackfynn as a .ttl file.
- CURATOR: Hand off image datasets to MBF Biosciences curators for segmentation assistance, spatial registration and conversion to SPARC approved formats and to transform the banner image.
- CURATOR: Hand off data If genetic or physiology data are present to the Auckland curation team to create appropriate data visualizations for those data types.
- CURATOR: Finalize the dataset within Blackfynn, adding the finalized description once data is aligned to the SPARC standards, annotated and sign off is received from the MBF Biosciences & Auckland team, adding license information and provisioning a DOI, if the data are to be published immediately.
- INVESTIGATOR: Final check by PI of dataset after curators sign off.
- INVESTIGATOR: Request dataset to be published
- CURATOR: Publishes the dataset (Principal Investigator), or allows it to be published automatically after the embargo period ends.

These steps can be viewed within the private data portal using the “dataset status”, a feature implemented in Blackfynn in December 2019. The steps that each dataset go through are formalized, numbered, and color coded (Fig. 11). Each label is associated with the party that is responsible for setting the particular status. Please note that the teams at MBF Biosciences and ABI are considered curators for this workflow. These teams are responsible for ensuring that SPARC data are aligned to common spatial frameworks, as described in the introduction.

**Figure 11:**
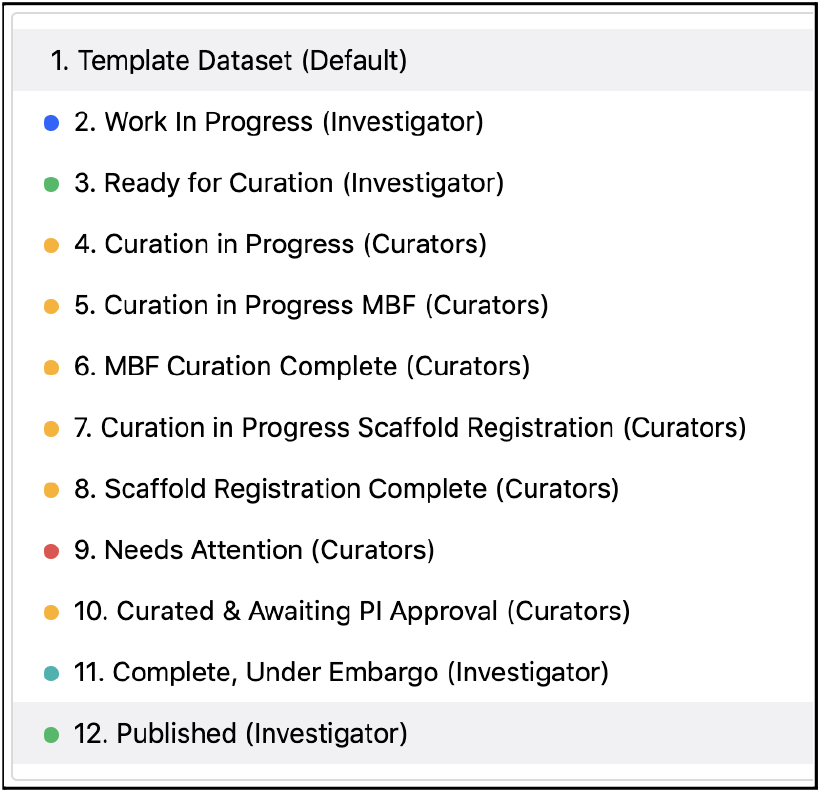
Ordered status types set by Investigators or set by Curators.

These steps are not necessarily performed in sequential order. For example, the image registration, conversion and segmentation performed by MBF Biosciences may be performed before the imaging files are uploaded to Blackfynn. Researchers do, however, create the necessary dataset descriptors in Blackfynn and often upload the necessary metadata files. This will mean that in some cases the order will go from 1-2-5-6(MBF Biosciences)-3-4(UCSD Curation).

Fig. 12 is a schematic representation of the workflow described above. It highlights how data is generated by individual investigators, curated by the Data Curation Team, and shared as an embargoed dataset with the SPARC Embargoed Data Sharing Group. It shows how the data is made available to the public over time.

**Figure 12:**
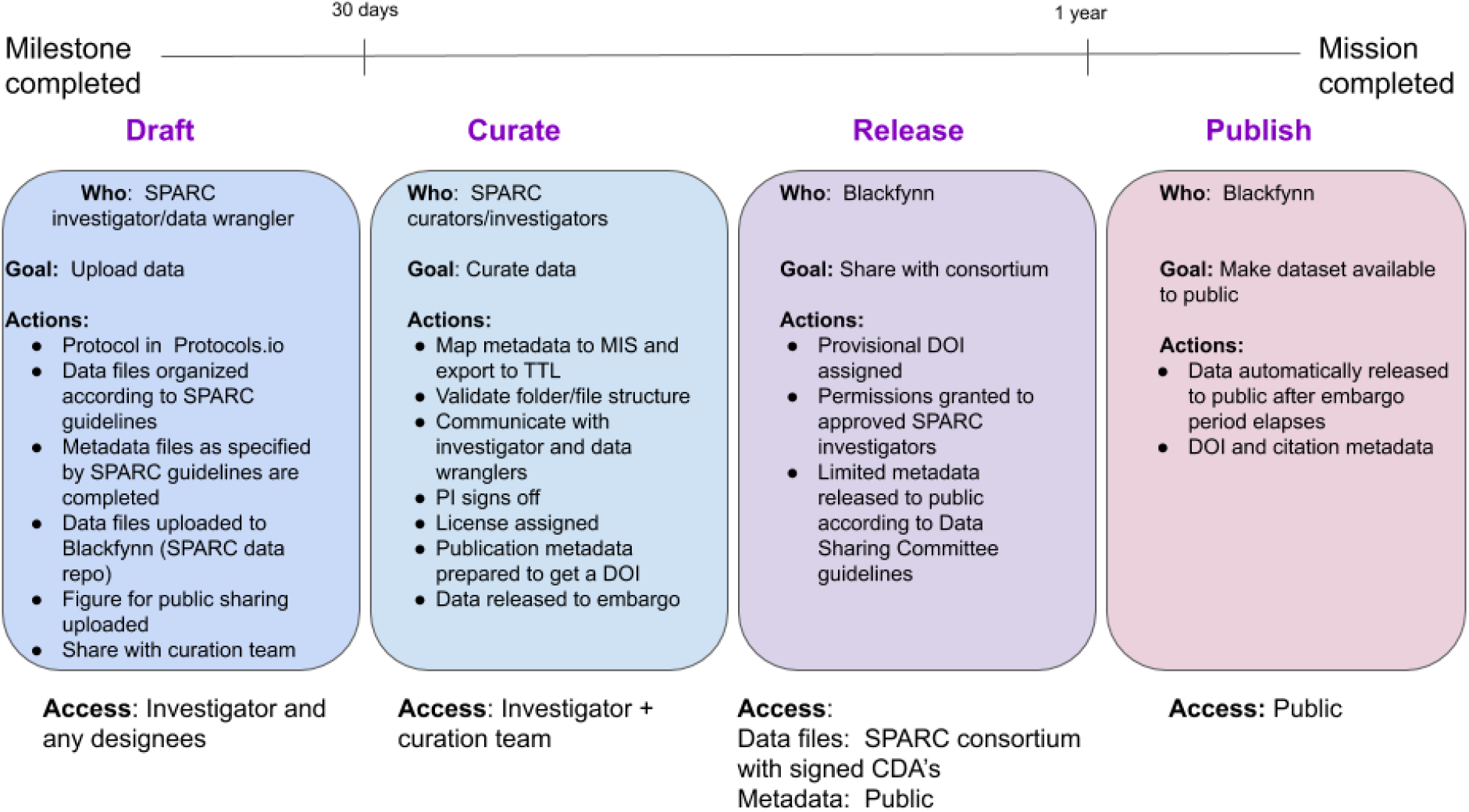
Overview of the entire submission-curation-publishing workflow.

## 8. Quality Control Metrics for SPARC Datasets

The SPARC curation team has developed a set of QC guidelines that are used to check for errors and to ensure consistency in the descriptions of SPARC data. SPARC datasets are checked by the curation team for the following:

1. They conform to the requirements of the SPARC Data Set Structure
2. The files are appropriately organized into the primary, derivative, docs, code, protocol folders
3. Manifests are included at each level of the folder tree and contain a sufficient description of the files present.
4. Title and description are clear, appropriate and detailed^15^.
5. If the data are part of a larger dataset, that relationships are specified
6. The species is appropriately identified and referred to consistently across protocol, dataset description and experimental data
7. All types of experimental data referred to in the abstract or protocol are contained in the dataset
8. All abbreviations used to describe the dataset across the different documents are defined
9. All experimental or sample groups referenced in the metadata are defined
10. All file types submitted conform to approved file types (upcoming)
11. s metadata standards are defined they are used appropriately

A checklist has been developed, which also includes questions that can be asked of the investigator^16^. Some of these checks will be incorporated into future versions of SODA.

## 9. Discussion

The establishment of the SDS has proven to be essential to curating the complex and large datasets submitted for SPARC. With a common structure, curation can take advantage of tools such as the validator to help with the curation process, thereby allowing them to focus on the scientific aspects of the dataset, e.g., is the description and protocol clear, rather than simple mechanistic tasks such as checking whether the number of subjects listed matches the number of subject folders. We realize that the SDS presents extra work for SPARC investigators, who must adapt their local lab practices to comply with a new structure. However, with the launch of SODA, investigators should find it easier to walk through the curation process. Finally, as the SPARC portal evolves, the user interface can take advantage of the regular structure to make it easier to browse SPARC data in a consistent manner.

In order for the SPARC project to meet its deliverables, the first round of standards needed to be implemented relatively quickly. The first public data released for SPARC occurred in July 2019 at the ISAN 2019 meeting. At that time, curators were curating to SDS 1.0, but many of the datasets released were demo datasets and were not fully structured. The SDS was revised in October of 2019 in response to the July release and through discussions with investigators. Data for the February 2020 release was curated to SDS 1.2.3. At that time, all of the original datasets were also recurated.

As the SDS continues to evolve - version 2.0 is scheduled to be released in spring of 2021 - we are not planning on recurating older data, as it would not be feasible to constantly revise the large number of datasets available through SPARC. We are, however, extracting larger amounts of structured information from these datasets, e.g., from the experimental protocols, and mapping it to the MIS, so some re-curation of metadata does occur. This information will be used to create more powerful and nuanced search across SPARC datasets and models.

Because of the Consortium’s variability in experimental methodology and primary data types, we are continuing to evaluate whether the SPARC Data Structure is sufficient for any investigator to unambiguously interpret the datasets from other research labs. On the technical side, we are examining this new structure’s ability to facilitate datasets to be exchanged and queried freely as well as understood by other scientists. For each use case such as simulation data or physiology data, we look at the relevant SPARC protocols and current results to determine the required parameters needed to understand the resulting data. We are using this information as a basis for formalizing modality-specific extensions to the SDS and MIS and to develop QC guidelines, as outlined in Section 8.

There are several additional areas where standardization will benefit SPARC. Within the next year, SPARC will also move to implement more consistent file formats for major data types, ensuring that SPARC data is available in non-proprietary formats. For example, all imaging data will have to be submitted as JPEG2000 and BioTIFF or in a format that can be converted to these formats. Guidelines for additional data types will be released in summer of 2021.

A third driver of standards in SPARC is the requirement to be interoperable with other data repositories, particularly those being created by the US BRAIN Initiative and other large brain projects around the world. The US BRAIN Initiative is investing in the creation of standards for major data types such as neuroimaging (BIDS^17^), neurophysiology (NWB^18^) and standards for 3D microscopy. These standards underlie the major archives established for BRAIN data: OpenNeuro, DANDI and the Brain Image Library, respectively. SPARC will be monitoring these standards for maturity and will create the means for SPARC data to be converted into these formats.

The establishment of standards for SPARC also underwent a governance change after the first data release. While in the first phase of the project, data standards were developed or recommended by the Data Standards Committee comprising SPARC investigators, after the first sets of data were released, responsibility for recommending and implementing new standards was shifted to the curation team, as they are most familiar with the breadth of SPARC data and the areas requiring standardization. The recommendations of the SPARC curation team are then put forward for review by the Data Standards Group and the SPARC community at large.

## Acknowledgements

We thank funding from NIH SPARC OT2OD030541, NIH SPARC OT2OD025308, and NIH SPARC OT2OD030213. The authors wish to thank Dr. Joost Wagenaar, Dr. Sue Tappan, the SPARC Data Standards Team and Ms. Ellen Goldstein for helpful input. 89/*

## Author contributions

A.B., J.G., A.P., T.G, G.P., M S-Z, and M.M. form the SPARC Curation Team and have all participated in the development of the SDS. B.P. is leading the development of SODA for SPARC. All have contributed to the writing and revision of this manuscript.

## Competing interest statement

AB, MM and JG have equity interest in SciCrunch.com, a tech start up out of UCSD that develops tools and services for reproducible science, including support for RRIDs. AB is the CEO of SciCrunch.com.

## Appendix Metadata specifications for SPARC datasets V1.2.3

**Table A1:**
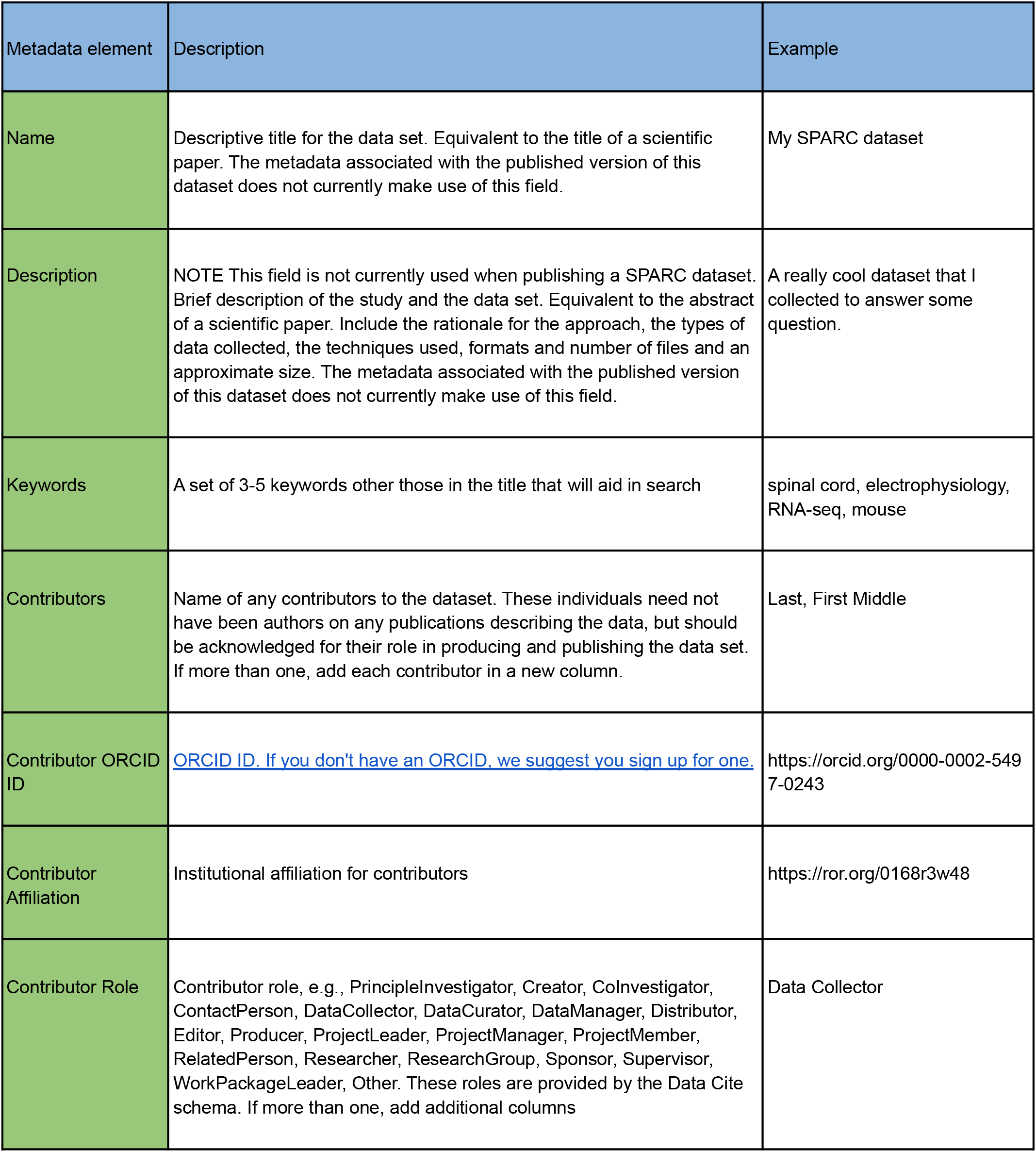
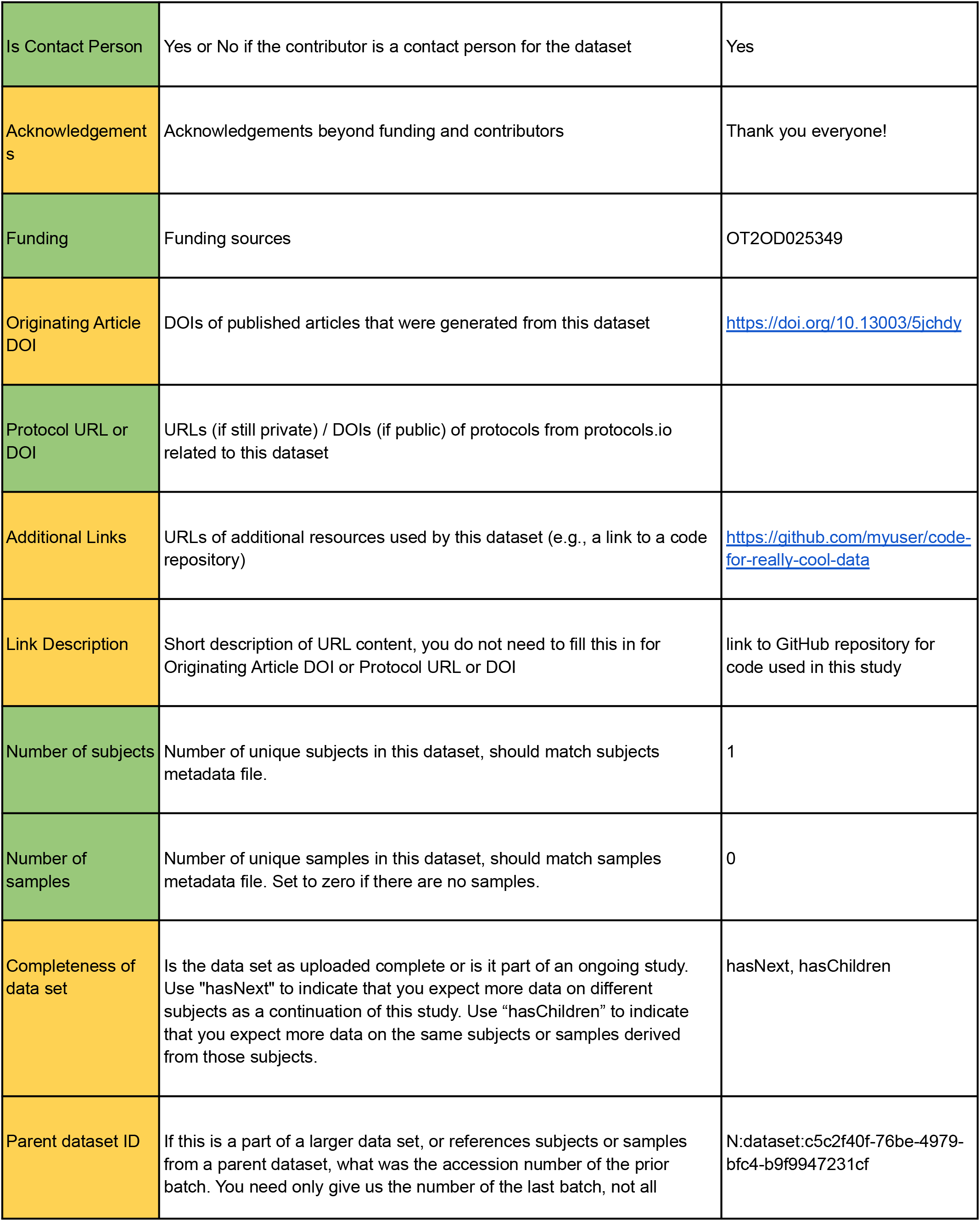
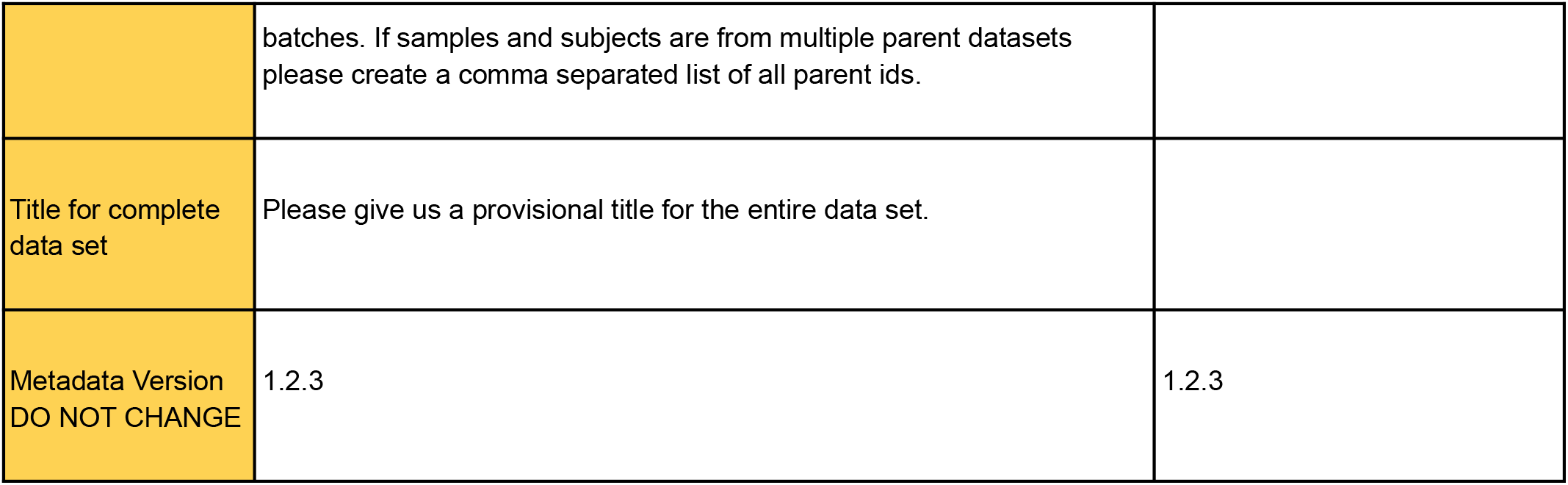
Descriptive metadata for V1.2.3. Required fields are highlighted in green while conditional fields (i.e., required if present) are highlighted in yellow.

**Table A2:**
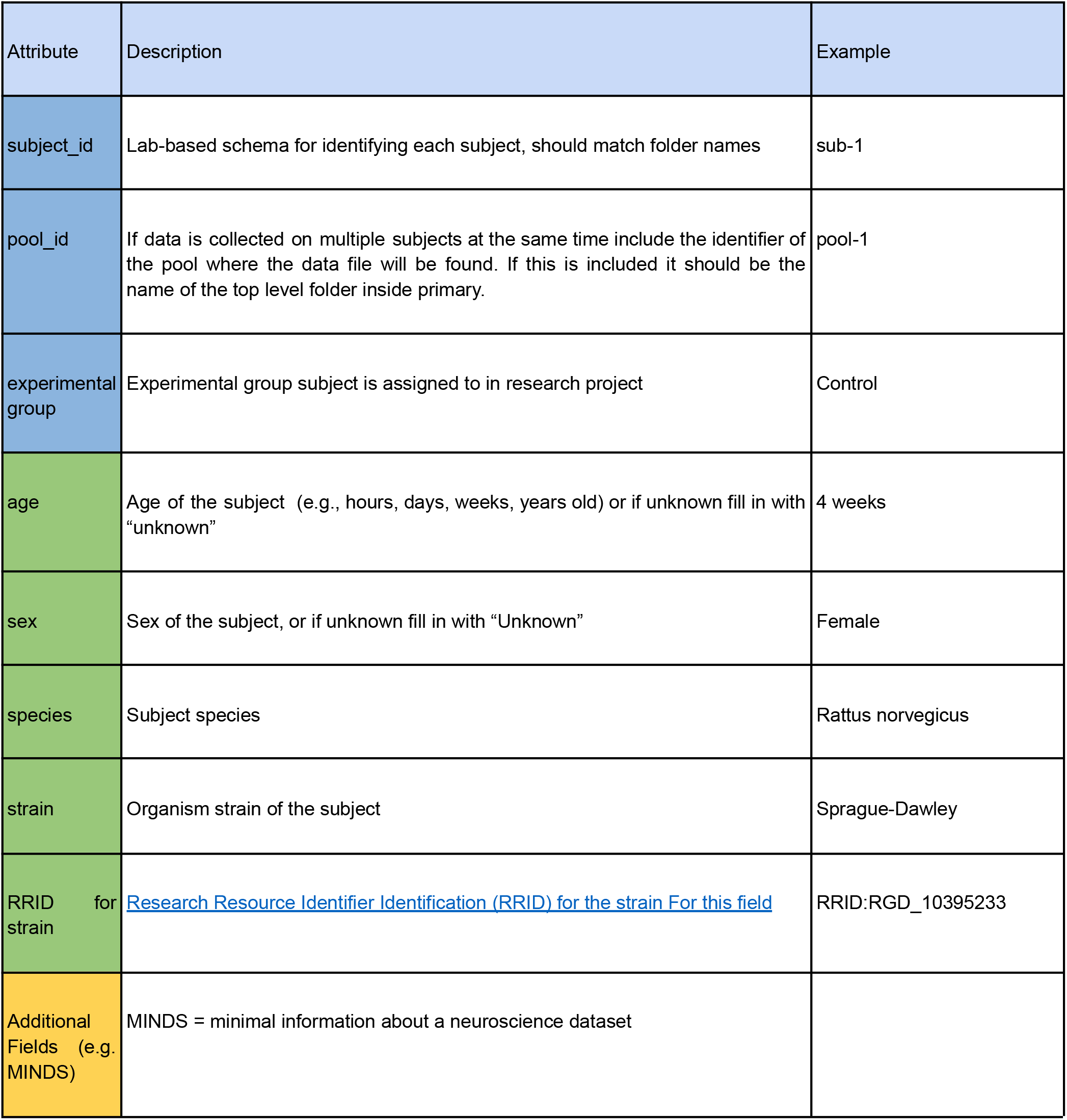
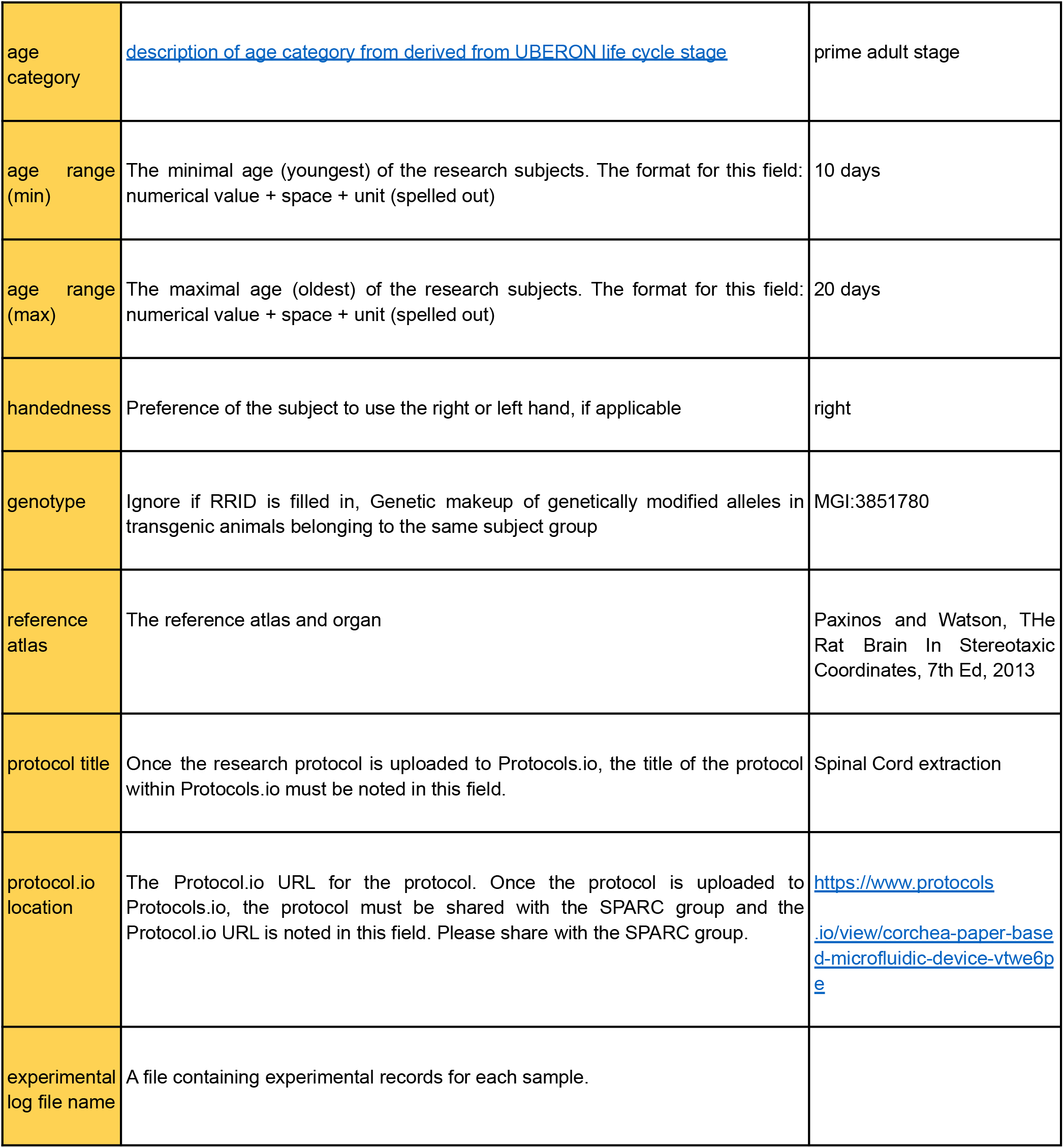
Subject metadata. Required fields are highlighted in green while recommended fields are highlighted in yellow. Blue fields are required (pool_id only if pooled subjects were used) and provide the necessary fields for providing provenance of subjects and subject pools within experiments.

**Table A3:**
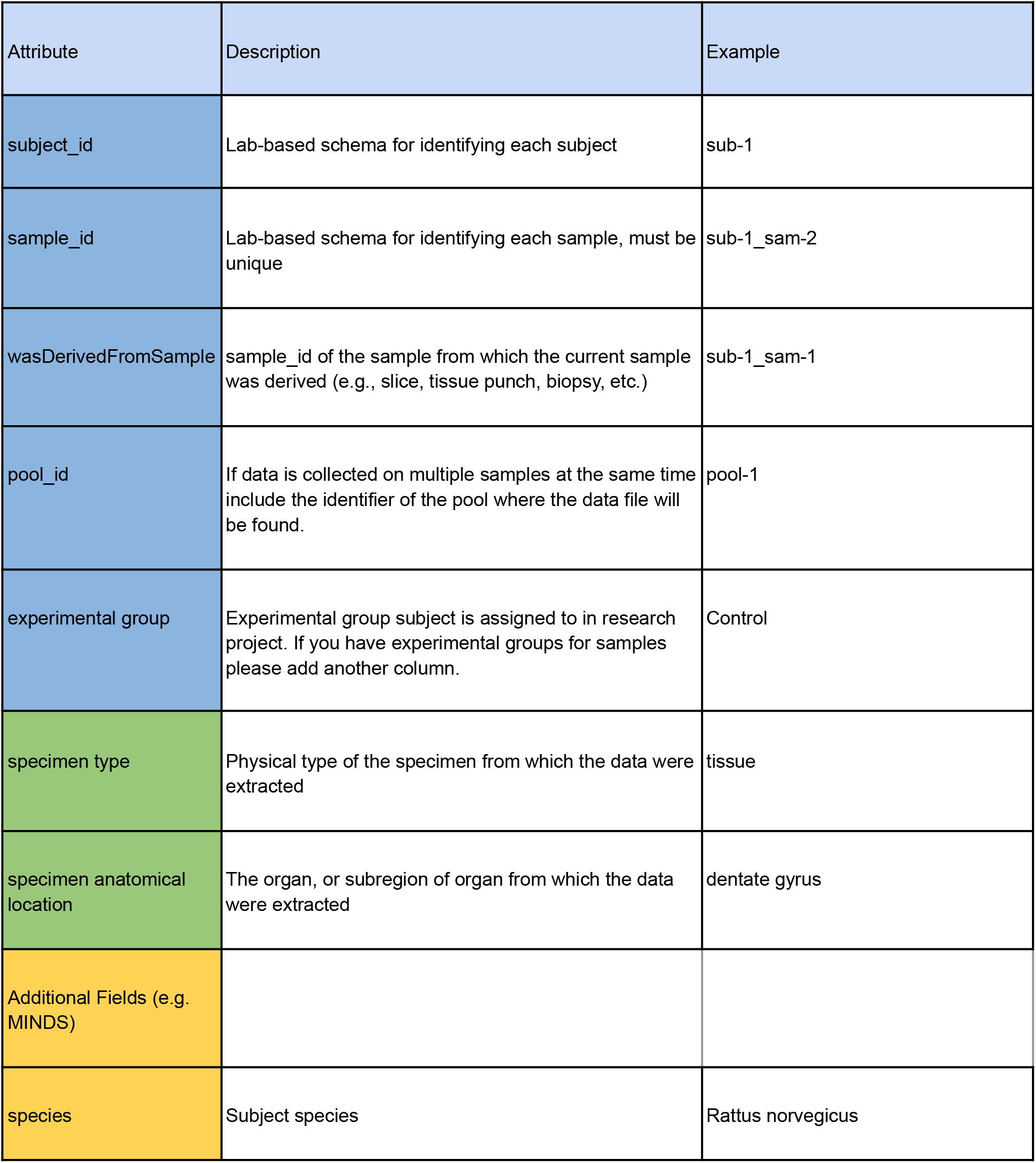
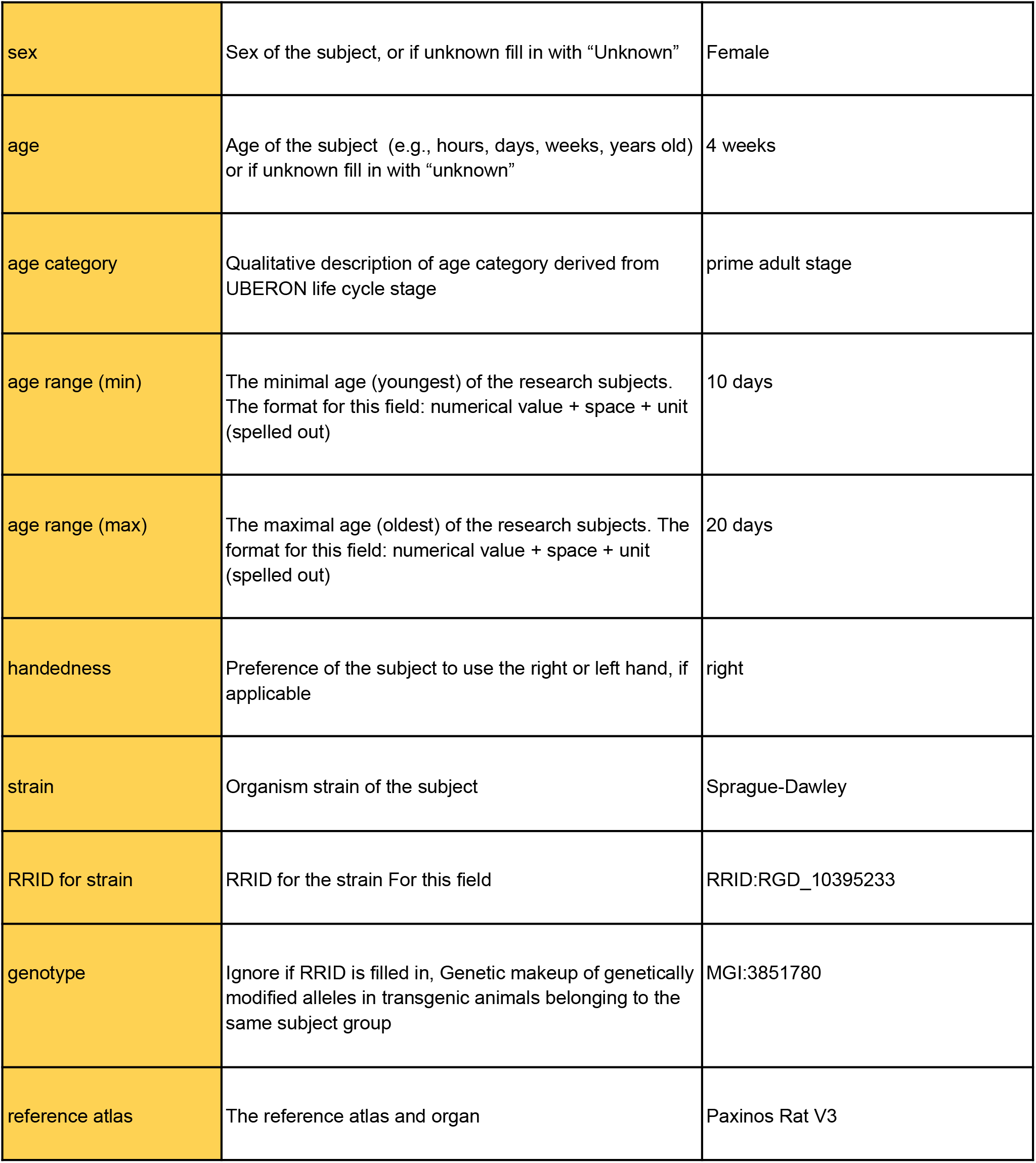
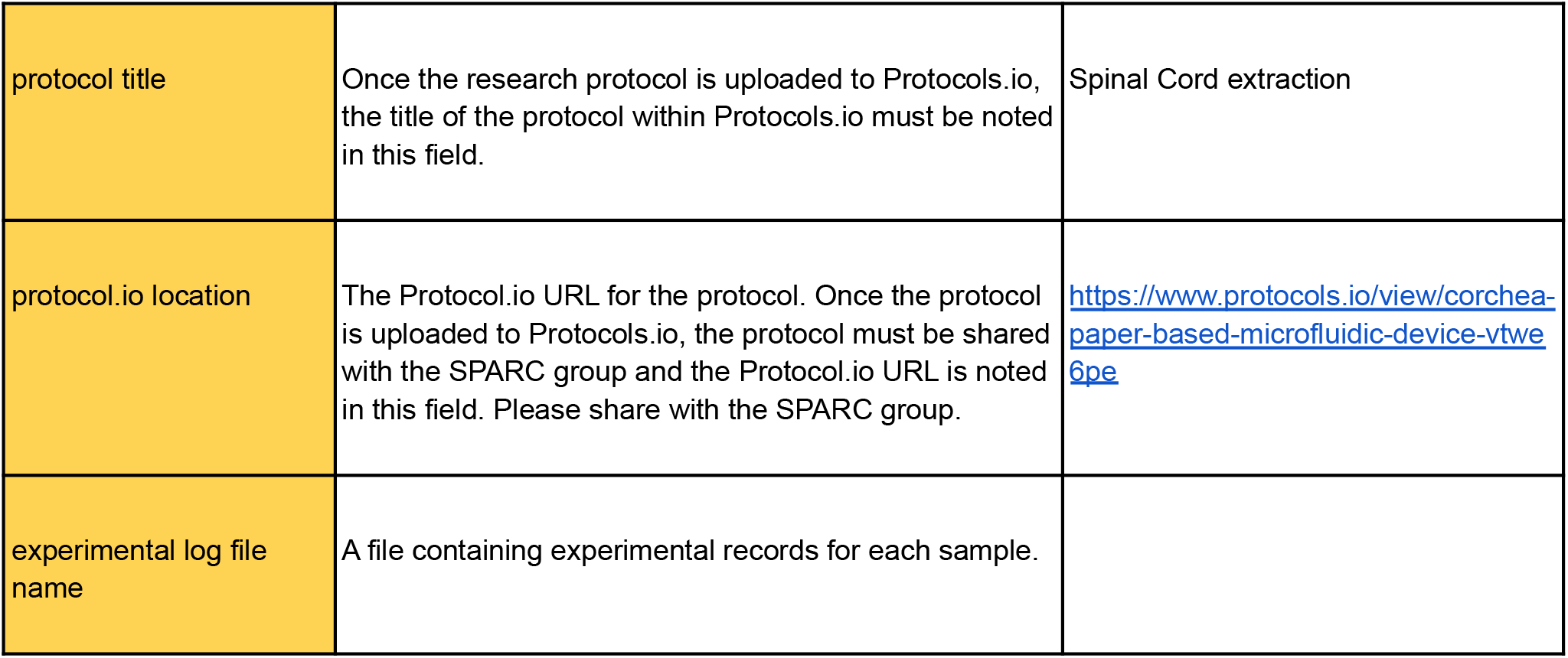
Sample metadata The color key is the same as for subjects (A2)

**Table A4:** Controlled vocabulary for experimental modes used in SPARC. These terms are in the process of being added to the NIFSTD ontology techniques branch. They are available in Interlex as children of experimental modalities: http://uri.interlex.org/base/ilx_0770148

| Term | Definition | NIFSTD |
| --- | --- | --- |
| behavioral approach | Study that induces and/or measures the behavior of the subject | http://uri.interlex.org/base/ilx_0739413 |
| cell morphology | Study that specifically seeks to understand the shape and structure of individual cells | http://uri.interlex.org/base/ilx_0739394 |
| cell population characterization | Study that measures biochemical, molecular and/or physiological characteristics of populations of cells as opposed to individual cells | http://uri.interlex.org/base/ilx_0739408 |
| computational modeling | Study that creates or characterizes computational models or simulations of other experimentally observed phenomena | http://uri.interlex.org/base/ilx_0739414 |
| connectivity | Study that maps or measures functional and/or anatomical connections between nerve cells and their targets or connections between populations of neurons in defined anatomical regions. | http://uri.interlex.org/base/ilx_0739393 |
| electrophysiology | Study that measures electrical impulses within an organism, cell or tissue or the effects of direct electrical stimulation | https://scicrunch.org/scicrunch/interlex/view/ilx_0741202 |
| epigenomics | Study that measures modifications of genetic material that affect transcription but do not alter the organism's DNA | http://uri.interlex.org/base/ilx_0741207 |
| expression | Study that measures or visualizes gene or protein expression within cells or tissues. Focuses on the gene. | http://uri.interlex.org/base/ilx_0739397 |
| expression characterization | Study that characterizes the cellular, anatomical, or morphological distribution of gene expression. Focuses on population. | http://uri.interlex.org/base/ilx_0739409 |
| genomics | Study that measures aspects related to the complete DNA genome of an organism | http://uri.interlex.org/base/ilx_0741204 |
| histology | Study that investigates the microscopic structure of tissues | http://uri.interlex.org/base/ilx_0739399 |
| microscopy | Study that primarily uses light or electron microscopic imaging | http://uri.interlex.org/base/ilx_0739404 |
| morphology | Study designed to determine the shape and structure of tissues and body parts | https://scicrunch.org/scicrunch/interlex/view/ilx_0739403 |
| multimodal approach | Study that employs multiple modalities in significant ways | http://uri.interlex.org/base/ilx_0739395 |
| physiology | Study that measures the function or behavior of organs and tissues in living systems. | http://uri.interlex.org/base/ilx_0739410 |
| radiology | Study that uses at least one of a variety of minimally invasive probes such as x-rays, ultrasound, or nuclear magnetic resonance signals to capture data about the internal structure of intact subjects. | https://scicrunch.org/scicrunch/interlex/view/ilx_0739390 |
| spatial transcriptomics | Study used to spatially resolve RNA-seq data, and thereby all mRNAs, in individual tissue sections (Wikipedia). | http://uri.interlex.org/base/ilx_0739396 |
| transcriptomics | Study that measures RNA transcription in the organism or cell population of interest | http://uri.interlex.org/base/ilx_0739406 |

## Notes

### Summary of Updates

We have addressed some typos and added so information that was missed in the previous version.

